# Conserved Human CpG Dinucleotides Identify Enriched Ageing-related Signals in Developmental Pathways and the Brain

**DOI:** 10.64898/2026.02.20.707043

**Authors:** Paraskevi Christofidou, Christopher G. Bell

## Abstract

The DNA methylome changes with age. This is observed as both random drift, but also consistent alterations within certain genomic loci enabling the construction of precise age predictors. However, the functionality of these ageing-related modifications remains largely undefined. The CpG dinucleotide is the principal sequence target for these epigenetic DNA marks in differentiated cells.

Here, for functional insight, we identified ultra-conserved CpGs (ucCpGs, n ∼167k) lacking observed sequence mutation in large-scale human whole genome datasets (>576k).

ucCpGs were enriched, as expected, in lowly-methylated CpG dense loci, due to methylated cytosine hypermutability. Additionally, ucCpGs demonstrated pathogenic evidence (CADD, ClinVar), and enrichment in four-fold degenerate sites, as well as within developmental and ageing-related gene families (AP-2, HOX-L, C2H2-ZNF, *etc*). Extreme ucCpG clusters (≥16 ucCpGs/kb) were enriched for brain-expressed genes, as well as developmental and ageing/mortality pathways. ucCpGs also strongly co-located within ageing-related Differentially Methylated Regions (age-DMRs), highlighting Clustered Protocadherin Gamma, as well as *HSPA2* and *LHFPL4* genes.

These findings further support that functional components of DNAm ageing are intertwined with developmental pathways.

## Introduction

In the human genome, DNA methylation (DNAm) of the cytosine base is essential for viability. This is clearly demonstrated by knockout studies of the DNA methyltransferase enzymes DNMT3A/B, the *de novo* ‘writers’ of this epigenetic modification^1,2^. Furthermore, disruption of epigenetic regulation, through genetic deletion of critical regulatory sequence carrying DNAm, can lead to severe developmental disorders^3,4^.

The CpG dinucleotide is the primary template for DNA modification in differentiated cells and therefore, represents a crucial genome-wide signalling module^5^. However, this palindromic two-base sequence is significantly depleted to approximately one-quarter of its expected occurrence in the human genome^6^. This loss arises because methylated cytosine (5mC) is prone to spontaneous deamination to thymine^7^, leading to a mutational rate ∼14-fold greater than other single nucleotide substitutions^8^.

DNAm is critical in enabling development and robust cellular identity, although, cell-specific DNA methylomes also change with age^9^, and represent a hallmark of ageing^10^. These changes occur in a quasi-stochastic fashion, with certain genomic contexts predisposed to specific directional shifts in DNAm^11^. For example, DNA hypermethylation of Polycomb Repressive Complex 2 (PRC2) target promoters and hypomethylation of solo-WCGW sites are recognised ageing-associated signatures^12–14^. The ability to measure DNAm at single CpG resolution, coupled with machine learning (ML) approaches, has enabled the development of highly accurate epigenetic ‘clocks’ that predict chronological and biological age^15^. These models have revealed strong epidemiological links between DNAm patterns and age-related morbidity and mortality^16^. However, deciphering all the contributory components summated in these ageing-related biomarkers is complex. These include, in commonly assessed peripheral blood, cell proportion changes due to biological processes such as immunoageing^17^. Furthermore, fascinating pan-mammalian DNAm analysis has revealed that a universal mammalian DNAm ageing clock can be constructed, and provided evidence that ageing is evolutionarily conserved and developmentally programmed^18^.

In this study, with the emergence of large-scale whole-genome sequencing (WGS) datasets, we have taken a human population genomics approach to assess CpG genetic conservation as a measure of functional importance. We have examined how these conserved loci intersect with ageing-related and tissue-specific changes. Genetic variation at methylated CpGs and fitness effects have been explored within coding regions previously, identifying highly deleterious sequence alterations^19^. Although current genomic biobanks lack comprehensive global representation – especially African ancestries – they nonetheless provide unprecedented power to identify conserved CpGs and infer their potential biological relevance.

## Results

### CpG variation in Whole Genome Sequencing Datasets

#### Ultra-conserved CpGs exhibit moderate overlap between gnomAD and UK Biobank

An analysis of ∼76k and ∼491k whole genomes from the gnomAD and UK Biobank (UKB) studies, respectively, was undertaken to determine the genome-wide landscape of variation at CpG sites. The UKB dataset is predominately of European ancestry whilst GnomAD includes greater population diversity (see Methods). As expected, the majority of CpGs were classified as variant CpGs (vCpGs) in both datasets (GnomAD vCpGs n=25,072,835; UKB vCpGs n=26,416,433) (Fig. 1A). After quality control for sequencing depth coverage and quality of variants (see Methods), we identified robust sets of ultra-conserved CpGs (ucCpGs) in GnomAD (n=1,020,349) and UKB (n=1,029,974). Notably, despite UKB being ∼6.5x larger, the number of ucCpGs was similar between datasets. This clearly reflects the broader demographic breadth of gnomAD, which captures greater global diversity, compared with the more genetically homogeneous UKB cohort. Furthermore, only ∼16% of the individual cohort ucCpGs were shared between the two datasets, resulting in a consensus set of 167,062 ucCpGs (Fig. 1B, Suppl. Data File 1). The genomic distribution of these ucCpGs across autosomes was generally uniform, although with some variation (Fig. 1C, Suppl. Table 3), *e.g*., chromosome 19 had the highest proportional ucCpG prevalence consistent with its known elevated gene density^20^.

**Fig 1:**
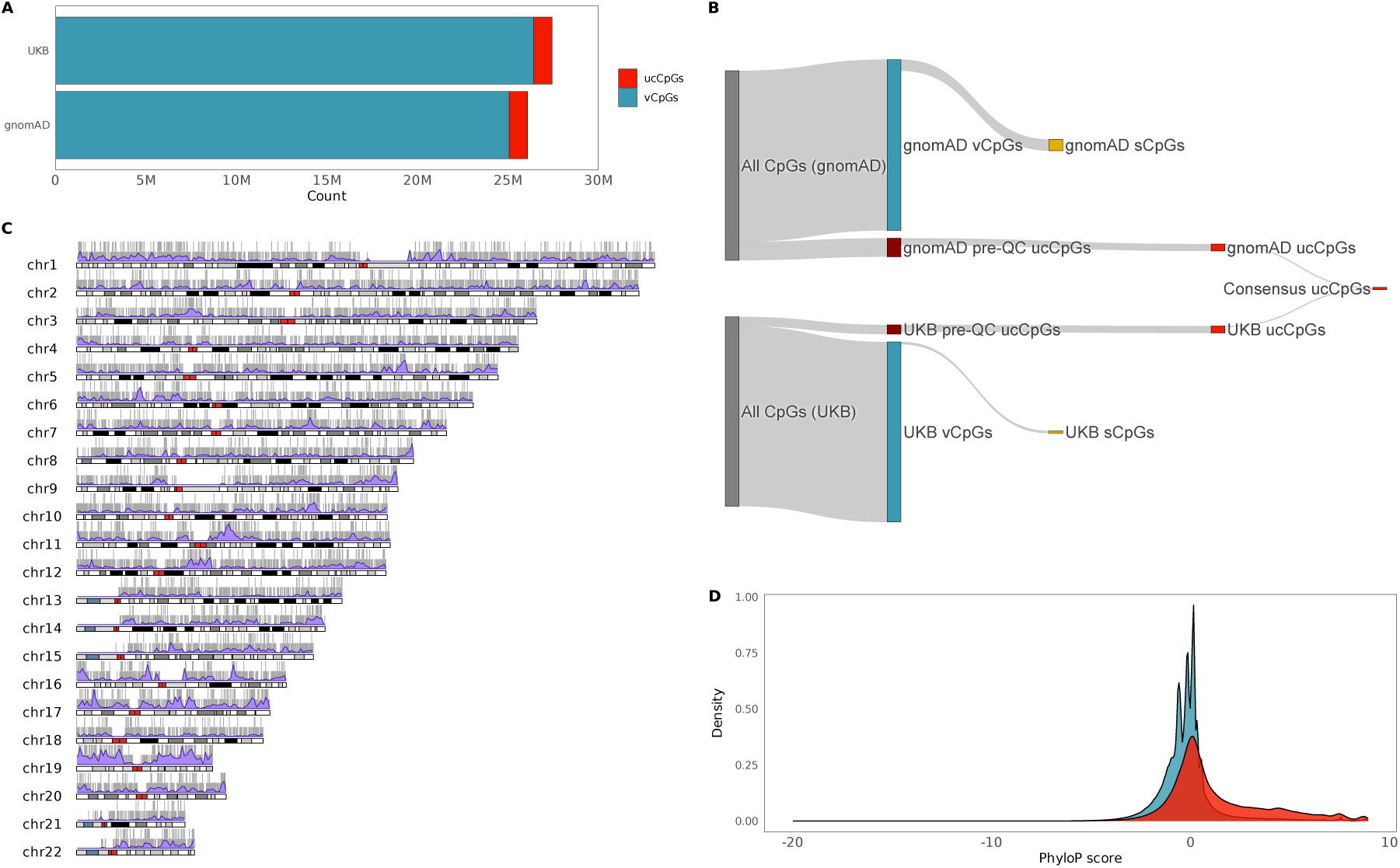
Ultra-conserved CpG (ucCpG) identification and distribution. **a**. Total number of post-QC variant CpGs (vCpGs) and ultra-conserved CpGs (ucCpGs) across gnomAD and UK 100 Biobank. **b**. Sankey plot illustrating the steps used to identify the consensus set of ucCpGs (sCpG: singleton CpG). **c**. Karyotype plot of overlapping ucCpGs. Ideogram represents cytogenetic banding patterns (dark bars: gene-poor; heterochromatic and light bars: gene-rich, euchromatic areas; red bars: centromeres; gaps: known assembly issues in GRCh38); ucCpGs distribution: solid grey lines, with overlapping ucCpGs stacked; ucCpGs density: purple, per 1Mb genomic windows. Higher peaks indicate regions with a greater number of ucCpGs, highlighting genome-wide regional enrichment patterns. **d**. Density plots show the distributions of average PhyloP scores for vCpGs (blue) and ucCpGs (red).

#### Ultra-conserved Human Population CpGs show expected Evolutionary Conservation

We assessed ucCpG evolutionary constraint via PhyloP scores (240 placental mammals)^21^. As expected, these strongly human sequence-conserved ucCpGs exhibited significantly higher PhyloP scores (Mean PhyloP: −0.21 vCpGs vs 1.26 ucCpGs; Wilcoxon p<2.2×10^−16^; Fig. 1D), indicative of population mutational constraint correlating with strong evolutionary constraint^22^.

### Ultra-conserved CpG enrichments

#### Ultra-conserved CpGs are Enriched in Proximal Regulatory Loci

We next examined the sequence-level distribution of the consensus set of ucCpGs around TSSs (Fig. 2A, Suppl. Table 4). Given that DNA methylation (DNAm) increases CpG mutability, ucCpGs would be anticipated to disproportionally reside in loci lacking germline DNAm, such as CpG Islands (CGIs).

**Fig 2:**
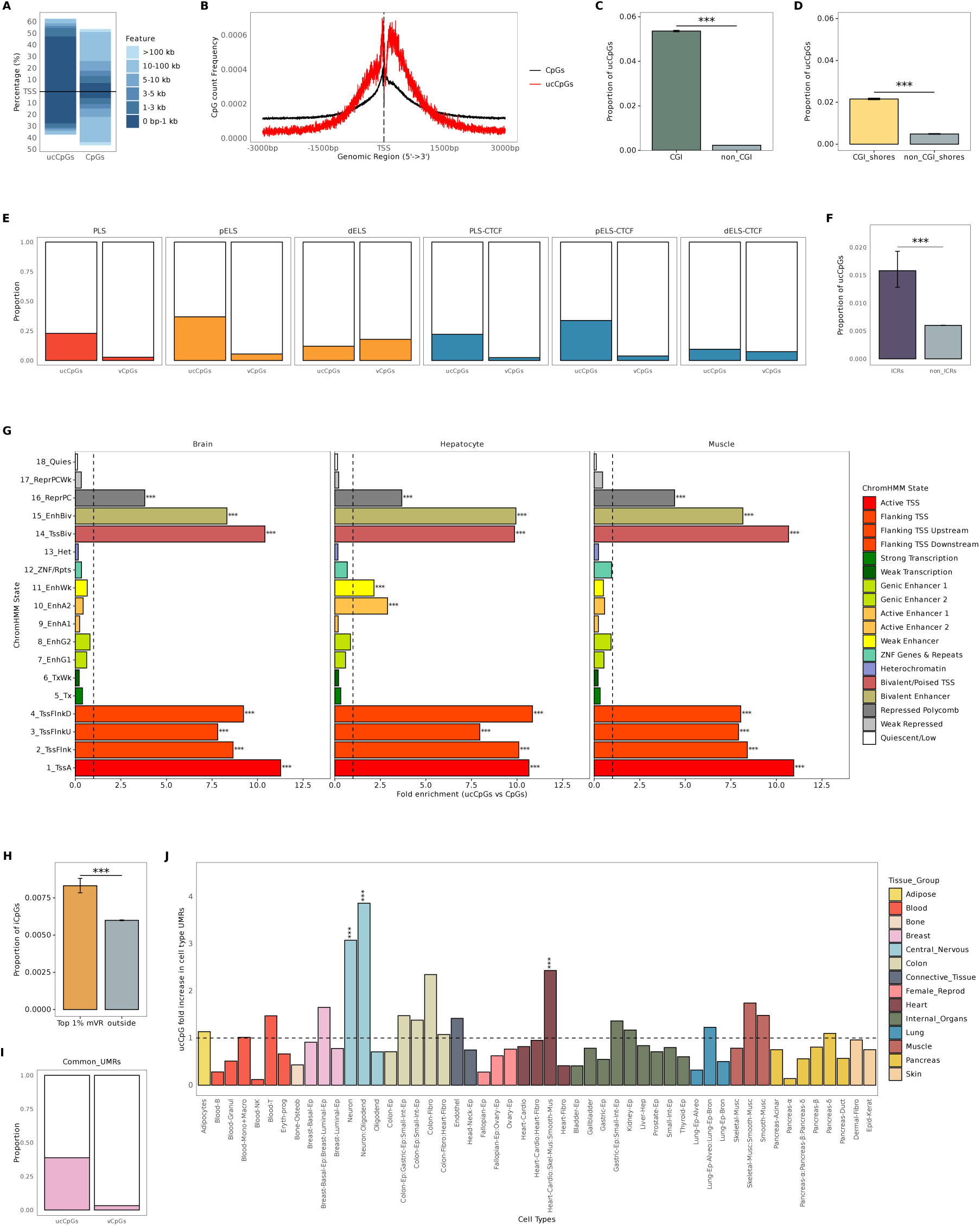
Enrichment analyses of ucCpGs. **a**. Distribution of ucCpGs and CpGs relative to TSS. **b**. Profiles of all CpG and ucCpG counts relative to TSS regions. **c**. ucCpGs enriched within CpG islands (CGI). **d**. ucCpGs enriched within CpG island shores (CGI_shores). **e**. ENCODE SCREEN cCRE: Promoters (PLS), proximal and distal enhancers (pELS/dELS) and CTCF-bound counterparts (All p<2.2×10^−16^). **f**. ucCpGs rare but still enriched within imprinting control regions (ICRs). **g**. ChromHMM state enrichment of ucCpGs across tissues. From DNAm atlas derived data^27^: h. ucCpGs within top 1% DNA methylation variable regions (mVR); i. ucCpGs enriched within Tissue Common Unmethylated Regions (UMRs); and j. ucCpG enrichment within cell-type specific Unmethylated Regions (UMRs). Vertical black line, point of neutral enrichment (y=1).

In line with this, the majority of ucCpGs are located within promoter regions (74.9% 0-1kb and 10.7% 1-3kb of the TSS) compared with only 24.2% of all CpGs (p<2.2×10^−16^).

ucCpG density gradually increases starting ∼1.5kb upstream of the TSS and extending up to ∼2kb downstream, indicative of co-locating CGIs (Fig. 2B). Notably, ucCpGs exhibited a pronounced enrichment peak approximately 30–40 bp upstream of the TSS (Peak: ucCpG= −37bp vs CpG= −1bp), and marked local depletion immediately downstream (Dip: ucCpG=86bp vs CpG=132bp). This pattern may reflect increased mutation associated with transcription initiation, potentially influenced by transcription factor (TF) binding, chromatin accessibility, directional or strand-specific mutation or repair, related to RNA Pol-II stalling and R-loop formation^23^. Consistent with this, recent work identified an appreciable uptick in CpG>TpG mutation ≤100bp downstream of the TSS^23^.

ucCpGs were strongly enriched within CGIs (OR:24.21, p<2.2×10^−16^, Fig. 2C, Suppl. Table 5). However, CGI shores (flanking CGIs ±2kb), regions that are often more dynamic in DNAm changes, particularly in response to environmental or developmental factors^24^, still resulted in an enriched signal (OR:4.51, p<2.2×10^−16^, Fig. 2D).

To further investigate the regulatory context of ucCpGs, we assessed their enrichment across six candidate *cis*-regulatory element (cCRE) categories as defined by ENCODE SCREEN^25^. This revealed a significant enrichment of ucCpGs (all p<2.2×10^−16^, Fig. 2E, Suppl. Table 6) within promoter-like sequences (PLS) and proximal enhancers (pELS) compared to vCpGs, as expected due to their CpG density preponderance. In contrast, low CpG density distal enhancers (dELS) showed significant depletion, suggesting less selective pressure to maintain CpG integrity at these loci, commonly captured as low methylation regions (LMRs)^26^. In contrast, strongly proximal and more subtly distal CTCF-bound loci were enriched.

#### Individual ucCpGs are rare but enriched within Imprinting Control Regions

We identified that ucCpGs were significantly enriched compared to the autosome-wide ucCpG/CpG ratio in critical imprinting control regions (ICRs, OR:2.66, p=1.54×10^−16^, Fig. 2F, Suppl. Table 7), where DNAm regulates parent-of-origin-specific gene expression. However, of the 68 well-characterised ICRs^28^ examined, they were only present in 37, and were rare, comprising only ∼1.6% of the total CpGs.

#### ucCpGs are enriched in bivalent/poised promoters

To identify chromatin-defined epigenomic functional enrichments, we compared the genomic distribution of ucCpGs with ChromHMM genome segmentation data in three developmental germ layer representative tissues: brain (ectoderm), muscle (mesoderm) and hepatocyte (endoderm). Overall, ucCpGs displayed highly similar chromatin state distributions across all three tissues. As expected, ucCpGs were highly enriched in promoter-associated states marking active and flanking transcription start sites (TSS) (TssA, TssFlnk, TssFlnkU and TssFlnkD) and significantly depleted in quiescent states (Quies) (Fig. 2G, all p<2.2×10^−16^, Suppl. Table 8). Beyond canonical promoters, ucCpGs were also significantly enriched in poised promoters found near bivalent TSS (TssBiv) and bivalent enhancers (EnhBiv) (all p<2.2×10^−16^). In addition, we observed significant enrichment of ucCpGs within Polycomb-Repressed (ReprPC) chromatin states (all p<2.2×10^−16^). This is consistent with the known role of Polycomb domains in silencing key developmental genes, which are often under strong evolutionary constraint. Furthermore, hyperconserved CpGs are known to underlie Polycomb-binding sites vital for embryonic development^29,30^.

#### ucCpGs are uncommon but enriched within highly cell-type variable DNA methylation loci

We next investigated the DNAm patterns of ucCpGs across diverse cell-types in the DNA methylation atlas^27^. This multi-tissue resource identified genomic regions showing the highest variability in DNAm across 39 cell types (top 1% variable DNAm blocks defined with ≥4 CpGs/block, ∼21k blocks). These highly cell-type discordant regions are indicative of cell-type-specific regulatory loci, such as enhancers. Although only a small proportion of ucCpGs were located within these blocks, they were still significantly enriched compared with their low total CpG background (OR:1.39, p<2.2×10^−16^; Fig. 2H, Suppl. Table 9). This indicates that, while uncommon, ucCpGs are preferentially retained in regions exhibiting dynamic methylation variation across cell types, consistent with a potential role in cell type specific regulatory architecture.

#### ucCpGs are common and enriched within pan-tissue unmethylated regions

Tissue-common unmethylated regions (UMRs) remain consistently unmethylated across multiple tissue types, and are predominantly associated with promoters and consequently CGIs. Consistent with our previous findings, ucCpGs were strongly enriched within UMRs identified from aggregating the cell-type specific data from the DNAm atlas (OR:19.25, p<2.2×10^−16^, Fig. 2I, Suppl. Table 10).

#### Neural cell-type specific unmethylated regions (UMRs) are enriched for ucCpGs

Whilst ucCpGs were not enriched within the overall set of cell-type specific UMRs, strong enrichments emerged when analysing the individual cell types (Fig. 2J, Suppl. Table 11). Specifically, we observed Bonferroni signficant enrichment within neuronal UMRs (OR:3.32, p=1.01×10^−12^), and in combined neuronal & oligodendrocyte progenitor UMRs (OR:4.24, p=1.75×10^−19^) but not within oligodendrocyte UMRs alone (OR:0.72, p=0.28). The multipotent progenitor of heart, cardiomyocyte, skeletal muscle and smooth muscle cells was also enriched (OR:2.57, p=2.91×10^−6^).

#### ucCpGs are enriched within Developmental and Ageing-related Gene Families

We explored the enrichment of ucCpG counts compared to the background number of CpG in HGNC gene families (see Methods). This identified 608 ucCpG-enriched gene families (p-adj<0.05, Suppl. Table 12), with 178 reaching Bonferroni significance (p<1.19×10^−5^). The top 25 results are displayed in Fig. 3A and identified gene families involved in developmental processes, disease mechanisms, and the ageing process. This included Nuclear Receptor Subfamily 2 Group F (*NR2F*), TFs intergral in organ and the central nervous system development (OR:3.78, p=2.38 ×10^−20^)^31^; the AP-2 TF family (*TFAP2A*-*E* OR:2.63, p=3.90×10^−17^), fundamental in embryonic and neurodevelopment^32–34^; and Class F frizzled G protein-coupled receptors (OR:3.39, p=1.51×10^−40^) critical in Wnt embryonic developmental pathways^35^. The *HOXL* class was also highlighted (OR:1.91, p=3.15×10^−38^) containing the *HOX* master regulator TFs of cell fate and body plan^36^.

**Fig 3:**
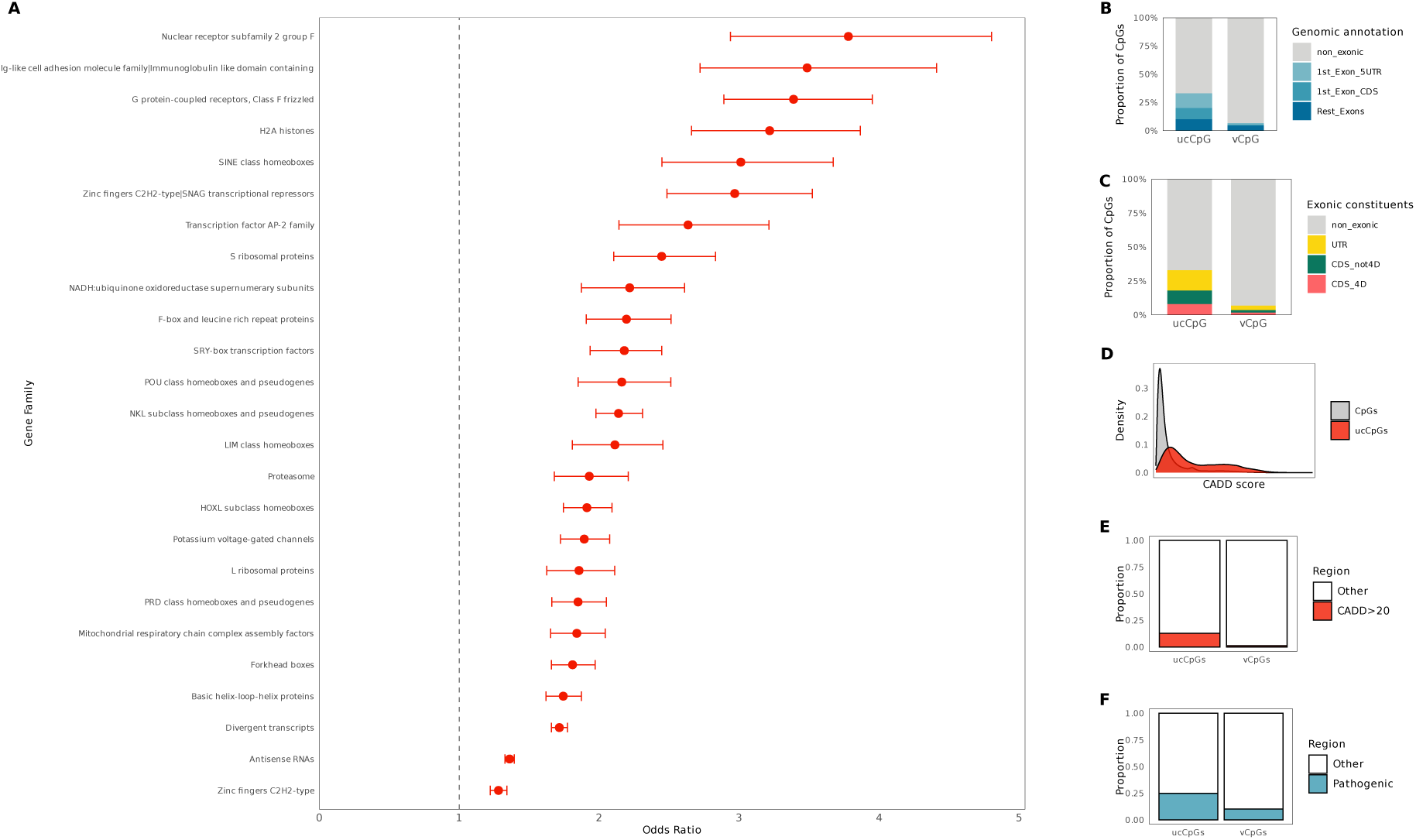
ucCpG enriched Gene Families, 4d sites, and pathogenic implications. **a**. Top 25 significant ucCpG enriched HGNC Gene families ranked by Odds Ratio. CpGs were annotated within defined gene promoters, exons & UTRs (hg38). **b**. ucCpGs enriched within 1^st^ exon 5’UTR and coding sequence (CDS). **c**. ucCpGs enriched within human 4-fold degenerate sites (4d). d. Density plots showing the distribution of mean CADD scores for all CpGs and ucCpGs. **e**. Proportion of ucCpGs enriched in CADD scores >20. f. Proportion of ucCpGs enriched in pathogenic ClinVar variants.

Gene families implicated in ageing included Histone 2A (OR:3.22, p=4.49×10^−27^) through the canonical^37^ and variant forms H2A.Z^38,39^ and H2A.J^40^; Basic helix-loop-helix proteins (bHLH) (OR:1.74, p=1.13×10^−44^), with intriguing evidence from *C. elegans* models^41^; and C2H2-type zinc finger (ZNF) protein TFs involved in the regulation of cellular senescence and ageing^42^ (OR:2.97, p=2.90×10^−27^). POU class homeobox genes are also enriched (OR:2.16, p=1.24×10^−19^, including *POU1F1*/*OCT4*) and have a significant role in development, but also ageing through stem cell pluripotency and cell identity maintenance^43^, as well as the Snell dwarf mutant *Pou1f1* mouse^44^. Interestingly, the bHLH and AP-2(*TFAP2*) genes mentioned above also show positive correlation with ageing in the universal DNAm mammalian analysis^18^, as well, the DNAm levels of *HOXL* subclass genes were associated in that study with maximum life span^18^.

#### ucCpGs are strongly enriched within 4-fold degenerate Mammalian Synonymous Sites

Almost one-third of ucCpGs overlap protein coding sequence (n=55,015, ∼32.9% total ucCpGs, GENCODE v48) and, therefore, this coding potential could be strongly contributing to their conservation. However, this is strongly skewed (>^2^/_3_) to the first exon (n=38,173, ∼22.8% total) due to ucCpGs residing within CGIs, moreover half of these are found within the 5’UTR (n=21,591, ∼12.9% total, Fig. 3B). In total ∼15.0% of all ucCpGs reside within either the 5’ or 3’ UTR (n=25,037).

Furthermore, of those ucCpGs within protein-coding sequence, we observed an extremely strong enrichment, compared to all CpGs, for human synonymous 4-fold degenerate (4D) sites (n=13,189, ∼7.9% of all ucCpGs, *cf*. 458,423 vCpGs, p<2.2×10^−16^, OR:5.08, Fig. 3C, Suppl. Table 13)^45^. These 4D sites are proposed to be restrained due to CpG-conditional mechanisms, including intragenic DNAm preventing spurious transcripts^46^, and specific overlapping TFBS^47^, as well as the epigenetic regulation of developmental genes, especially those of the *HOX* family^45^. Also, contributing to high GC content aids the recognition of ‘native’ transcripts^45^.

#### A small proportion of ucCpGs are human-specific CpGs

ucCpGs show strong mammalian conservation, therefore, we did not expect a strong human-specific ucCpG signal. Intersecting with a set of ∼1.19M human-specific CpGs identified through 6-primate comparison^48^, we identified only 1,271 human-specific (hs) ucCpGs (∼0.76%). Of note, these hs-ucCpGs overlap three human-accelerated regions (HARs: HAR1/2, HAR106, HAR108)^49^ and 13 human ancestor quickly evolved regions (HAQERs) enriched for neurodevelopmental enhancers^50^ (Suppl. Table 14). These include the HAQER CpG beacons (*DPP10*[HAQER0261], *CHL1*[HAQER0297], *EMILIN2*[HAQER0223], and again *HAR1*[HAQER035]) where position-selection on GC-biased gene conversion has contributed to human-specific regulation^51^.

#### ucCpG mutation is associated with increased Pathogenicity

We evaluated ucCpG deleteriousness and clinical significance using CADD scores^52^ and ClinVar^53^. On average, ucCpGs showed substantially higher CADD scores compared to all CpGs (Mean CADD: 10.02 ucCpGs vs 3.33 CpGs; Wilcoxon p<2.2×10^−16^), indicating stronger functional constraint (Fig. 3D). Consistently, ucCpGs were markedly enriched in regions with high predicted deleteriousness (CADD> 20; 12.98% ucCpGs vs 1.41% CpGs, OR:10.94, p<2.2×10⁻¹⁶, Fig. 3E). ClinVar pathogenic variants were significantly enriched at ucCpG sites (24.9%) compared to all CpGs (10.5%; OR=2.9, p<2.2×10^⁻16^, Fig. 3F).

### Identification of extreme ucCpG clusters

#### Statistically Significant ucCpG clusters were observed with ≥16 ucCpGs/kb

To explore the genomic distribution of ucCpGs, we constructed ucCpG-centered 1kb windows (±499bp per ucCpG; n=167,062; see Methods/Suppl. Fig. 2). The ucCpG/kb density was predominately low (median:5, IQ range:3-8), so that even in CpG-rich regions, ucCpGs remain relatively rare (Fig. 4A).

**Fig 4:**
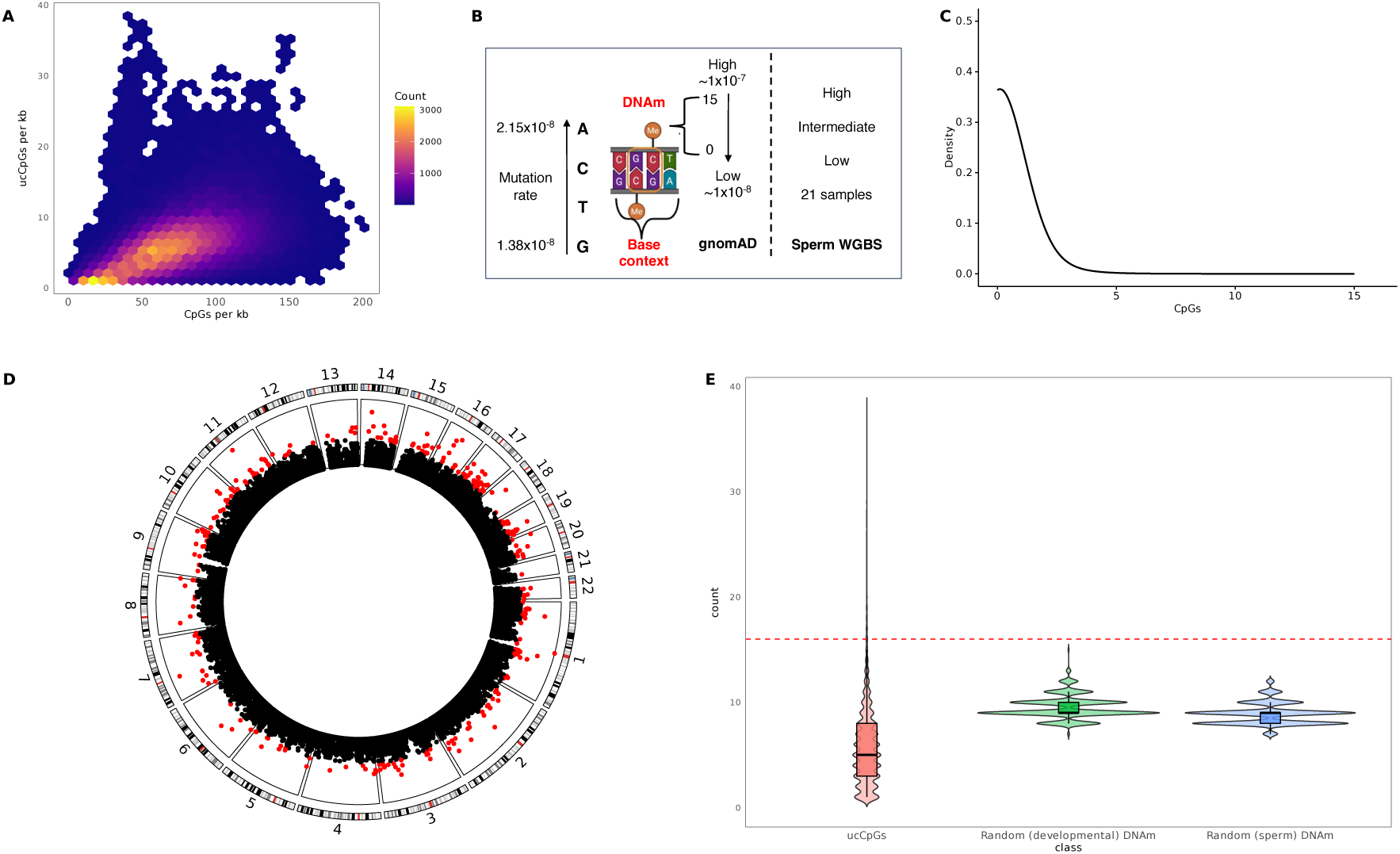
ucCpG cluster analysis. **a**. Smooth scatter plot of the distribution of 167,062 1kb windows based on their CpG (x-axis) and ucCpG count (y-axis). Colour scale gradient indicates the local density of windows (darker = higher). **b**. Schematic representation of steps used to calculate mutational likelihood. **c**. Plot depicting the distribution of ucCpG clusters across 1,000 permutations. d. Circular plot illustrating the distribution of ucCpG/kb clusters across the autosomal genome (red: ≥16ucCpG/kb). **e**. Violin plots illustrating the distribution of observed ucCpG clusters and that simulated data of random ucCpGs (green: GnomAD developmental dataset; blue: sperm dataset) never exceeded ≥16 ucCpGs/kb.

To account for their overlapping nature, the ucCpG windows were then merged into 30,270 contiguous unique ucCpG clusters (see Methods). Among these, 37.0% were ‘solo-ucCpGs’ containing only one ucCpG, while 37.1% contained ≥5 ucCpGs and 9.6% contained ≥10 ucCpGs ucCpG clusters were uniformly distributed across autosomal chromosomes (Fig. 4D, Suppl. Table 15).

To evaluate the statistical likelihood of high-density ucCpG clusters, we performed 1,000 mutability-weighted permutations. This approach accounted for the mutational likelihood of each CpG within its surrounding base context and its developmental/germline DNA methylation level^54^ (Fig. 4B, see Methods). 167,062 randomly selected CpGs, representing the ucCpG set, were selected from our total autosomal set of CpGs (∼27.7M) for each of the 1,000 permutations, and the simulated data never exceeded ≥16 ucCpGs/kb (Figs. 4C, 4E). Using this threshold, 311 extreme ucCpG clusters in the observed data (red) were classified as statistically significant (empirical p<1×10^−3^; Fig. 4D). To evaluate the robustness of this threshold, we repeated the 1,000 permutations with our mutation weighting calculated using DNAm data from a differing sperm methylome dataset^55^, and no simulated window exceeded >12 ucCpGs/kb.

To further identify whether natural groupings exist among ucCpG clusters, we applied an unsupervised machine learning approach using Gaussian mixture models (GMM, see Methods). The model was fitted on z-scored counts from ucCpG containing 1kb windows. In the 167k ucCpG windows, the GMM identified only four clusters (Suppl. Fig. 3), with the highest (≥13) largely overlapping with permutation-defined results, reinforcing the presence of biologically distinct, extreme ucCpG clusters. After filtering out loci overlapping common structural variants from recent global long-read analysis^56^ (see Methods), 298 extreme ucCpG clusters remained (Suppl. Table 16).

#### Extreme ucCpG cluster-associated genes are strongly enriched for Cerebral Cortex Expression and Loss-of-Function Intolerance

The 298 enriched ucCpG clusters comprise 8,135 individual ucCpGs, with the majority (71.6%) positioned within ≤1kb of the TSS (Fig. 5A). These ucCpGs also exhibited markedly higher conservation (via PhyloP) than both all CpGs and total ucCpG set (Fig. 5B).

**Fig 5:**
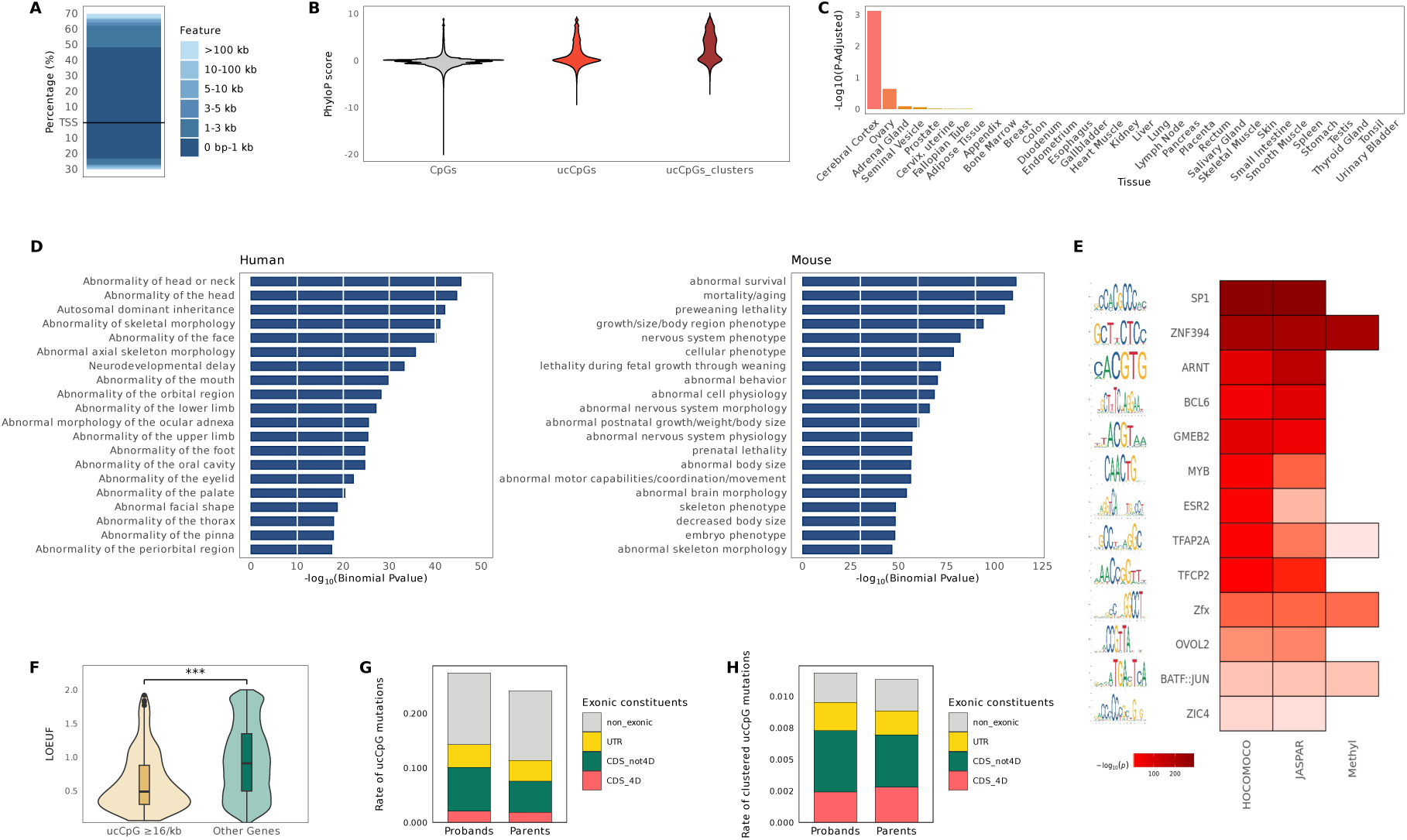
Gene Set, Transcription Factor & Rare Disease Mutation Enrichment. **a**. Distance of ucCpGs within extreme clusters relative to TSS. b. PhyloP scores across all CpGs, ucCpGs and ucCpGs within extreme clusters (≥16 ucCpGs). **c**. TissueEnrich results for ucCpG extreme cluster genes (data from Human Protein Atlas)^57^. **d**. Top 20 Gene Set Enrichment results from functional annotation and GO enrichment analysis of ucCpGs via GREAT. False discovery rate (p<0.05) adjusted binomial p-values are shown in the plot. e. Transcription factor motif enrichment analysis (-log10 evalues; gradient light to dark red with increasing significance) for motifs derived from HOCOMOCO, JASPAR, and DNAm sensitive motif predictions^58^. **f**. LOEUF (Loss-of-function Observed/Expected Upper bound Fraction^59^) scores measuring gene tolerance to mutation (lower scores indicating high intolerance and strong constraint) for ucCpG cluster genes versus other genes. g. Rate of mutation and distribution of exonic, UTR and 4d site mutated ucCpGs in Genomics England Rare Disease (RD) parents and probands **h**. Rate of mutation and distribution of exonic, UTR and 4d site mutated ucCpGs within ucCpG extreme clusters in RD parents and probands.

The TSS of 272 genes was located within ±5kb of these extreme ucCpG clusters (Suppl. Table 17). Examining tissue-specific expression patterns, we found these genes to be significantly enriched for cerebral cortex expression, the key brain region in complex cognitive functions (41/272 genes, log_2_ fold=1.97; Hypergeometric Adjusted p=7.56×10^−4^, Fig. 5C, Suppl. Table 18).

These genes were also significantly enriched for intolerance to loss-of-function (LoF) mutations, as measured by LOEUF scores (Fig. 5F). Low LOEUF values indicate high LoF intolerance and gene essentiality (Mean LOEUF: 0.63 ucCpG cluster genes vs 0.95 other protein-coding; Wilcoxon p<2.2×10^−16^).

#### Gene Set Enrichment of Extreme ucCpG Clusters implicate Development, Ageing, and Mortality associated Pathways

We perfomed region based Gene Set Enrichment for the 298 extreme ucCpG clusters via GREAT (Fig. 5D, Suppl. Tables 19-20). For human phenotypes, this revealed a significant enrichment for morphological and developmental phenotypes, including ‘Abnormality of head or neck’ (Bionomial p=2.75×10⁻^46^), ‘Abnormality of skeletal morphology’ (p=1.09×10⁻^41^), and ‘Neurodevelopmental delay’ (p=6.35×10⁻^34^). Additional enrichments included ‘Autosomal dominant inheritance’ (p=9.22×10⁻^43^) and multiple craniofacial and limb patterning defects, highlighting the developmental specificity of these loci.

Consistent with this, the corresponding mouse knockout orthologs showed exceptionally strong enrichment for phenotypes associated with survival, growth and nervous system function, including ‘Abnormal survival’ (p=7.74×10⁻^112^), ‘Mortality/ageing’ (p=5.21×10⁻^110^), ‘Preweaning lethality’ (p=8.10×10⁻^106^) and ‘Nervous system phenotype’ (p=1.12×10⁻^82^). These enrichments are similar to those identified by Lu *et al*. in the mammalian DNAm ageing analysis^18^. Together, these findings suggest that extreme ucCpG clusters demarcate genomic loci under strong purifying selection, preferentially marking genes essential for developmental integrity, neuronal function, and organismal survival across mammals, but also an intriguing overlap with ageing-related genes.

#### Transcription Factor Motif Enrichment highlights critical Developmental Regulators

We next investigated whether ucCpGs within extreme ucCpG clusters were enriched for regulatory transcription factor (TF) binding motifs (±10 bp). Motif analysis using JASPAR and HOCOMOCO collections revealed highly significant enrichments across both datasets (Suppl. Table 21). 14 TFs were consistently identified across both datasets (Fig. 5E). Among the overlapping TFs, four factors: SP/KLF-like, ZN502/SMCA5/ZN394, Zfx and TFAP2A, were also enriched in methylation sensitive TFs^58^.

An interesting observation was the enrichment of motifs for *TFAP2A* (E=4×10^−15^), a member of the AP-2 TF gene family, a critical developmental regulator in embryonic and neurodevelopment^32,60^, as we previously identified the AP-2 TF gene family itself to be ucCpG-enriched (Fig. 3A). Interestingly, *TFAP2A* DNAm was correlated with both ageing and lifespan in the mammalian ageing analyses^18,61^.

#### ucCpG and ucCpG clusters are enriched for mutations in a Developmental Disease enriched human cohort

We extended our analysis to the Genomics England (GE) rare developmental disease cohort (∼62.7k genomes: ∼29.6k probands & ∼33.1k parents), where we examined the mutation rate at ucCpG sites. Of the set of ∼167k ucCpGs, 15.5% (n=25,957) were mutated, with a higher rate uniquely within probands (8,085, ∼0.27/genome) compared to parents (7,962, ∼0.24/genome, hypergeometric p=6.38×10^−21^, Fig. 5G, 9,910 were in both) suggesting a potential pathogenic relevance for a subset. The signal did show evidence of being contributed to by ucCpGs overlapping CDS non-4d, but this did not explain all of the increase (Fig 5G, Suppl. Table 22). Examining only the ucCpGs within extreme clusters a slight non-significant increase was still observed in the probands versus parents (1.19×10^−2^/genome *cf*. 1.13×10^−2^, Fig. 5H, Suppl. Table 22).

### Ageing-related ultra-conserved CpGs

#### ucCpGs are enriched within mitotic DNAm ‘clocks’

Having identified overlaps with previous ageing-related findings, we next focused directly on our ucCpG dataset to interrogate the intriguing nature of the ageing DNA methylome.

Firstly, we contrasted the ratio of ucCpGs to CpGs for 1^st^-generation chronological DNAm clocks (Horvath^62^; Hannum et al.^63^; Horvath Skin & Blood^64^; & Zhang et al.^65^) in comparison to that proportion for their specific array background (Fig. 6A and Suppl. Table 22). For example, only 23 CpGs of the total 353 from the original Horvath clock are ultra-conserved and were not significantly enriched, considering the background set of 1,534 ucCpGs on the ∼25.9k 27k-450k common array CpGs that this clock was constructed from. Neither were the rest of these chronological clocks ucCpG-enriched. We then compared the ratios of two phenotypic clocks, the 2^nd^-generation PhenoAge^66^ and the 3^rd^-generation pace of ageing DunedinPACE measure^67^. Again, ucCpGs were present but not strongly statistically enriched, *e.g*., there was a non-Bonferroni significant enrichment for PhenoAge: 26 of 513 CpGs compared to a background of ∼13.5k ucCpGs on the 450k array (p=4.26×10^−3^).

**Fig 6:**
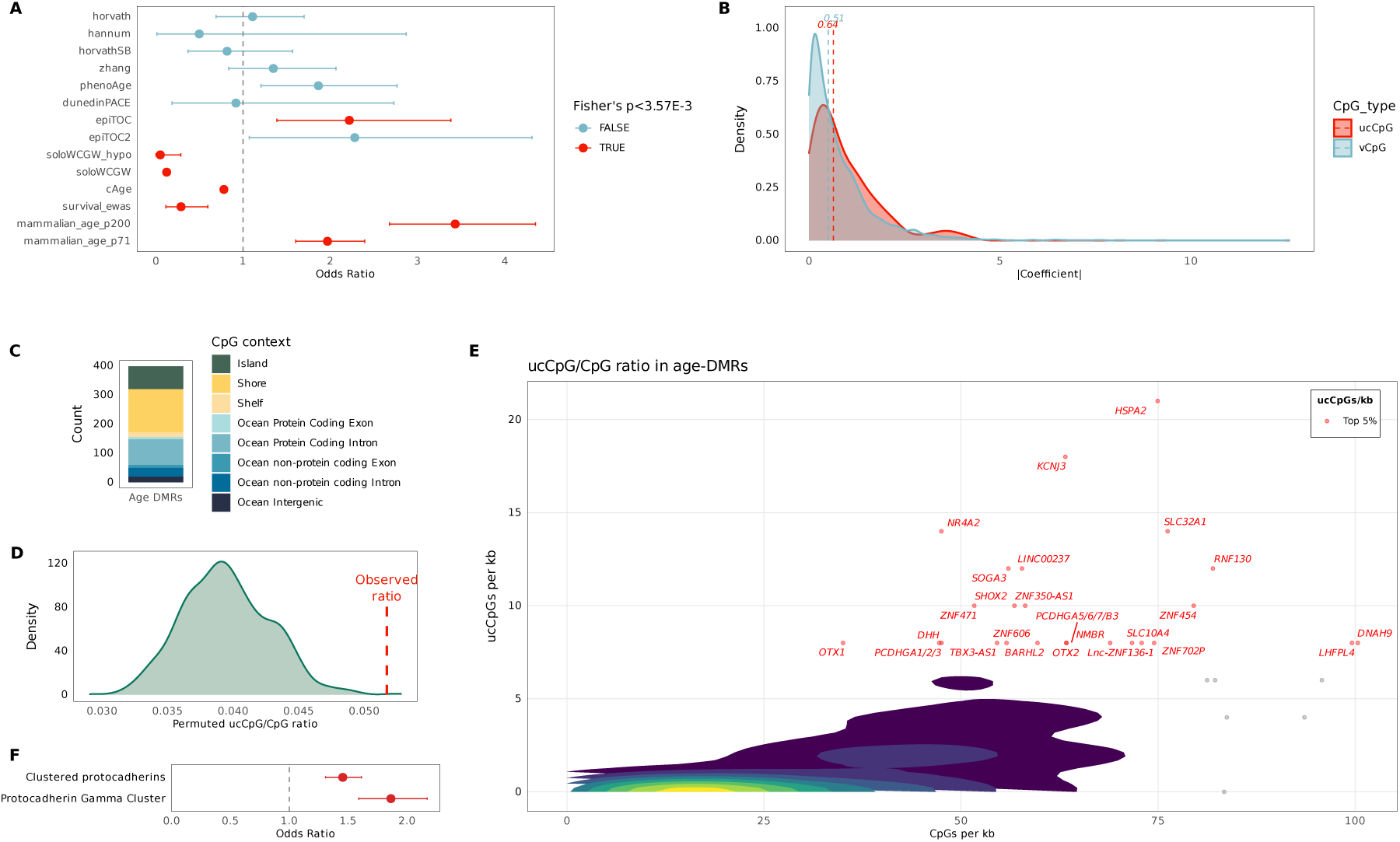
Age-related ucCpG analysis. **a**. ucCpG enrichment within DNAm ‘clocks’ (1^st^ generation: Horvath, Hannum et al., Horvath Skin & Blood, Zhang et al.; 2^nd^ Generation: PhenoAge; 3^rd^ Generation: DunedinPACE; Mitotic: EpiTOC, EpiTOC2, SoloWCGW-solo), total set of soloWCGW, cAge (Chronological Age predictor), Survival EWAS results, Mammalian DNAm ageing EWAS results (p<1×10^−200^ and p<1×10^−72^). **b**. Density plot of Chronological age CpGs linear coefficients by ucCpG (red) versus variant (vCpG: blue) status. **c**. CpG functionally context (CpG island, shore ±2kb, shelf ±2-4kb, and ocean by coding overlap) of the age-DMRs used for ucCpG age-DMR permutation. **d**. Permutation results of these age-DMR ucCpG overlaps. Observed age-DMR ucCpG/CpG result=0.052, permutation mean 0.039, range=0.029 - 0.053, Empirical p value=0.002. **e**. Smoothscatter plot of ucCpG/CpG ratio within the age-DMRs. The individual top 5% age-DMR results for ucCpG/kb (∼6.3% DMRs due to equal values) are indicated in red with their nearest gene (within +/-10k). **f**. HGNC gene family ucCpG/CpG enrichment for Clustered Protocadherins and the Protocadherin Gamma subfamily.

We then examined the two mitotic DNAm clocks specifically designed to capture cellular turnover, EpiTOC1^68^ and EpiTOC2^69^. These clocks are constructed from ageing-related promoter CpGs of Polycomb group target genes that are hypomethylated in multiple different fetal tissues. Through their measure of cell division, a positive acceleration has been associated with precancerous and cancer tissue risk^15^. Both clocks showed a positive enrichment for ucCpGs, with EpiTOC1 being Bonferroni significant (OR: 2.22, p=8.45×10^−4^, Fig. 6A).

We then turned from selected clock CpGs to broader ageing-related datasets. Examining the ∼13% of CpGs (∼3.7 million) recognised due to their sparse nature and specifically termed ‘soloWCGW’, due to a lack of neighbouring CpGs within ±35bp as well as being flanked by weak bases (A or T)^14^. These CpGs, found predominately within late-replicating regions as well as partially methylated domains (PMDs), are commonly observed as a group to lose DNAm stochastically with age^11^. ucCpGs are significantly depleted within this soloWCGW set, with only ∼3.1k ucCpGs (OR:0.12, p<2.2×10^−16^). This clearly fits with soloWCGWs being predominately hypermethyated and, therefore, hypermutable. Likewise, a mitotic ‘clock’ subset of only 678 of these CpGs represented on the 450k/EPIC arrays (soloWCGW hypoClock) was also significantly depleted of ucCpGs^69^.

#### ucCpGs are depleted within chronological age-associated CpGs, but a subset drives higher coefficients in chronological age predictors

Next, we interrogated the ageing results from one of the largest DNAm array datasets – the Generation Scotland analysis of ∼18k blood-derived DNAm arrays. This study constructed a chronological age predictor (cAge;^16^) by first identifying 99,828 chronological age-associated Differentially Methylated Cytosines (DMCs) from a curated array total of 752,695 CpGs. Only 1,626 of these cAge DMCs are ucCpGs, and, therefore, were strongly depleted (OR:0.78, p=5.40×10^−22^). This was consistent with our prior results from the 1^st^ generation chronological clocks. Furthermore, a survival EWAS in this dataset identified 1,182 significant CpGs, which was also depleted of ucCpGs (p=1.05×10^−4^). This latter result implies that ucCpGs are possibly less likely to occur in those CpGs that are influenced by environmental factors. To test this further, we looked at the overlap between ucCpGs and a smoking meta-analysis of ∼11k individuals^70^. This did indeed show a significant depletion with only 1,103 ucCpGs represented in the ∼65.8k smoking associated CpGs (OR:0.88, p=2.80×10^−5^, Suppl. Fig. 24).

However, whilst ucCpGs were in general depleted in cAge DMCs, it was interesting to note that when examining the subset of 2,275 CpGs employed in the constructed linear cAge predictor, the ucCpGs present here had a stronger absolute coefficient contributing to the prediction (see also quadratic results, Suppl. Fig 4). The median coefficient for ucCpGs was 0.64 versus 0.51 for vCpGs (Wilcoxon p=2.39×10^−2^, Fig. 6B). This effect was strongly driven by four ucCpGs with absolute coefficients >2.5: cg14911690 (*PBX4*); cg19401340 (*PPM1E*); cg08197797 (*PERP*); cg11220950 (*SYNGR3*). Therefore, whilst in totality ucCpGs are reduced in these chronological CpGs, there is some evidence that a small subset of ucCpGs disproportionality contribute to this signal.

#### ucCpGs are strongly enriched within Pan Mammalian ageing-related CpGs

We next examined the pan-mammalian ageing results from the specifically designed mammalian DNAm array^18^. CpGs selected for their strong mammalian conservation would be expected to also show strong conservation across human populations. The 1,214 ucCpGs (∼3.24%) of the 37,488 mammalian array CpGs were indeed enriched compared to their autosomal CpG fraction (∼0.60%). However, ucCpGs are even more enriched in the set of mammalian CpGs that show ageing-related changes (Fig. 6A). For top 2k mammalian age-DMP results (p<1×10^−71^), 118 are ucCpGs (5.9%, OR:1.97, p=3.08×10^−10^). This enrichment is even stronger for those top mammalian ageing CpGs exceeding p<1×10^−200^, with 82 ucCpGs of these 832 CpGs (∼9.86%, OR:3.43, p=4.83×10^−19^).

#### ucCpGs are also enriched within whole genome peripheral blood age-DMRs

We next explored results from a whole genome ageing-related DNAm analysis. These 398 age-Differentially Methylated Regions (age-DMRs) were derived from MeDIP-seq in peripheral blood with results defined within 500bp windows^71^. These age-DMRs possess a very strong enrichment for ucCpGs (∼5.17% ucCpGs, OR:9.04, p<2.2×10^−16^) but as both MeDIP-seq DMRs and ucCpGs are enriched for CpG dense regions, we performed additional analyses to further assess significance. Firstly, we performed 1,000X permutations by sampling windows of the exact size and CpG density of the age-DMRs and determining their ucCpG count (see Methods). This remained highly significant (Suppl. Fig. 5, observed ucCpGs in age-DMRs=428, versus permutation mean 207.2, range=139-287, Empirical p<1×10^−3^). To further consider CpG functionality, we performed an additional 1,000X permutation, randomly sampling the genome to match the CpG-defined context of the age-DMRs, i.e., CpG islands, shores, shelves, or ocean regions. Furthermore, we split the ocean regions into those that overlapped protein coding and non-protein coding exons and introns, and finally intergenic regions (Fig. 6C). This permutation was also significant (Fig. 6D, observed ucCpG to CpG ratio in age-DMRs=0.052, permutation ratio mean 0.039, range=0.029-0.053, Empirical p=0.002).

#### ucCpG enriched ageing DMRs highlight HSPA2, LHFP4, and the Protocadherin Gamma Gene family

To delineate the major drivers of this age-DMR ucCpG enrichment, we identified the top 5% age-DMRs for ucCpG to CpG ratio (Fig. 6E, Suppl. Table 25) indicated by their co-located or nearest gene (±10kb). This identified *HSPA2* as the strongest result with an ucCpG/CpG ratio of 21/74 within this 1kb age-DMR, strongly contributed to by 10 CpGs overlying 4d sites (Suppl. Fig 7). This small ∼2.5kb one exon gene encodes the Heat Shock Protein family A member 2, which shows high expression in testes but also multiple brain regions (Suppl. Fig 6). This gene is involved in protein and enzyme binding impacting multiple signaling pathways^72^. This hypermethylating age-DMR, initially identified in the context of DNAm ageing within an atrial fibrillation-associated GWAS locus^73^, also gains further support from a recent brain ONT long-read analysis that found a similar increasing DNAm with age (β=0.186^74^). Additionally, *HSPA2* possesses a consistent ageing-related decrease in expression (4^th^ strongest) across multiple human tissues in GTEx data for the RelativeAge^+^ module in the Mammalian analysis^75^. It was of interest to see that the *LHFPL4* gene was also highlighted that contains two CpGs that were the top and 3^rd^ result in the mammalian ageing EWAS (Suppl. Fig 8). This DMR is within the densest CpG region (∼20% CpG).

Two loci represented Protocadherin Gamma gene clusters were in this top group (*PCDHGA1*/*A2*/*A3* and *PCDHGA5*/*A6*/*A7*/*B3*) (Suppl. Fig 9). These protocadherin genes were recently identified in a multi-tissue DNAm pre-print from Jacques *et al*. as pan-tissue drivers of ageing^76^. In fact, whilst this gene family was not in the top 25, Clustered Protocadherin genes were significant in our gene family analysis (OR:1.45, p=1.47×10^−11^, Supp. Table 12). Furthermore, as the evidence is specific for the Gamma cluster, we selected the genes in this sub-family and identified that this was even more significant (Fig. 6F, OR:1.86, p=2.88×10^−13^). Additionally, the ONT long-read brain analysis, discussed above, identified that Protocadherin (*PCDH*) cluster promoters were among the strongest age predictors^74^.

## Discussion

This analysis into CpG functionality has been enabled by the unprecedented ability to interrogate greater than half a million human whole genome sequences, including >576k from GnomAD and UKB, plus an additional disease-related cohort of ∼63k from Genomics England.

We identified ∼167k ultra-conserved CpGs (ucCpGs). These strongly conserved dinucleotides were enriched to reside within developmental and ageing-related gene families, including *HOXL*, AP-2 (*TFAP2*), *C2H2-ZNF*, Histone 2A, *NR2F*, *FZD* (GPR-F), *POU*/*OCT*, and *bHLH*. This is consistent with prior work that developmentally critical Polycomb-bound genes are hypomethylated and subsequently hypomutated to maintain critical epigenetic regulation^29,30,45,47,77^. Furthermore, this supports the universal mammalian DNAm analysis where correlations with age were identified for many of these families, including bHLH & AP-2 (*TFAP2*) as well as an association with maximum life span for *HOXL* DNAm levels^18^. We additionally identified a neuronal UMR cell-type specific enrichment of ucCpGs from DNAm atlas data^27^.

Whilst groups of CpGs may work together to maintain regulatory function, synthetic methylation-silencing systems indicated that certain CpGs may have disproportional functional effects^78^. Supporting the potential significant functionality of ucCpGs, we identified increased CADD and ClinVar scores, as well as mutation rate within the Genomics England disease probands. This proposes that there may be clinical utility in exploring the ucCpG set further in developmental disorders. Exonic overlap revealed a very strong enrichment for CpG maintanence within synonymous 4d sites, indicative of the important epigenetic role these dinucleotides play in developmental and polycomb target genes^29,30,45^.

Using mutational weighting as well as ML we identified extreme ucCpG clusters with a conservative threshold (≥16 ucCpG/kb). These clusters were enriched in the promoters of brain-expressed genes as well as those critical genes more intolerant to mutation. These results have parallels with previous findings that increasing CpG density as well as PRC2 element EZH2 bound-promoters are associated with increasingly mutation intolerance genes^79^. Gene set enrichment of these regions identified developmental abnormalities, nervous system, and ageing/mortality pathways, mirroring those pathways identified in the Mammalian DNAm ageing analysis^18^. TFBS enrichments, including DNAm sensitive TFs, identified a number of TF groupings, including the developmentally important *TFAP2A*, where both the binding motifs and the gene family itself are enriched for ucCpGs, indicative of a coordinated epigenomic network^80^.

Ageing-related DNAm analysis revealed an enrichment for mitotic clocks, which are targeting PRC-binding developmental genes. Interestingly, a depletion of ucCpGs was shown within environmentally responsive loci (tobacco-related). Within genome-wide age-DMRs, *HSPA2* makes an intriguing candidate to follow up due to its wide signalling brief^72^, brain findings^74^, as well as pan-tissue reduced expression in the mammalian RelativeAge^+^ module^61^. *LHFPL4* is also clearly of interest with additional support from our analyses following up its top Mammalian DNAm ageing EWAS result^18^. The ucCpG-enriched Clustered Protocadherin Gamma (*PCDHG*) genes also have accruing evidence of importance in ageing having been recently identified as pan-tissue drivers of ageing^76^. Blood and brain DNAm of *PCDHG* were intriguingly recently identified to be influenced by the *KAP1*/*TRIM28* deSUMOlator *SENP7*^81^.

Clearly expanding our WGS analysis into more diverse populations would be beneficial and reduce the current ucCpG set^82^. However, even with this limitation, the dataset so far has identified intriguing functional enrichments in these strongly conserved loci. Furthermore, earlier analysis within coding synonymous and germline methylated sites indicates a core global invariant CpG (iCpG) subset does exist^19^. Ultimately only mutations never seen at CpGs as samples grow in size, would be those iCpGs that are embryonically lethal.

In conclusion, through human population sequence analysis we have identified fuctionally-enriched ucCpGs. These findings integrated with ageing-related changes support evidence that components of DNAm ageing are intertwined with developmental pathways and critical developmental genes. Further work to explore these developmental, brain, and ageing-related loci through functional epigenomic manipulation in appropriate cell model systems will be potentially revealing.

## Methods

### Human Genome CpG Dinucleotides

The reference set of CpG dinucleotide locations for human genome build GRCh38 (autosomes only) was ascertained from the RaMWAS R package^83^ (https://github.com/andreyshabalin/ramwas). This set comprised a total of 27,789,753 CpGs that passed QC (https://bioconductor.org/packages/release/bioc/vignettes/ramwas/inst/doc/RW3_BAM_QCs.html). These CpGs were identified by combining reference genome sequence data with SNV information (MAF > 0.01) from all super populations (European, African, Ad-Mixed American, East Asian and South Asian) included in the 1000 Genomes Project (Phase3). CpGs with unreliable coverage estimates due to poor alignment were excluded in this set.

### Whole Genome sequencing human samples

Genetic variation information derived from Illumina Short-Read Whole Genome Sequencing (WGS) on 566,855 individuals was obtained from two available sources: gnomAD^54^, and UK Biobank^84^. These datasets were interrogated for their allelic frequency at CpG dinucleotides genome-wide.

#### GnomAD

We utilized data from the Genome Aggregation Database (gnomAD), (https://gnomad.broadinstitute.org/data#v4) which comprises high-quality WGS from diverse populations, all mapped to the GRCh38 reference sequence. gnomAD v4.1.0 includes whole genomes from 76,215 individuals, aggregated from multiple sequencing projects, with rigorous quality control to remove low-confidence variants, related individuals, and individuals with severe paediatric disease. The genetic ancestry composition was distributed across the following groups: Europeans (n=5,316 Finnish and 34,025 non-Finnish: 51.62%), Admixed Americans (n=7,657: 10.05%), African/African American (n=20,805: 27.30%), Ashkenazi Jewish (n=1,736: 2.28%), Amish (n=456: 0.60%), East Asian (n=2,598: 3.41%), Middle Eastern (n=147: 0.19%), South Asian (n=2,417: 3.17%) and remaining (n=1,058: 1.39%). Variant calls include single nucleotide variants (SNVs) and small insertions/deletions (Indels). Population allele frequencies, variant annotations, and coverage metrics are available through gnomAD (https://gnomad.broadinstitute.org/data#v4). We downloaded the available VCF files for all autosomes and cleaned them based on allele frequency for each alternative allele (AF) being AF>0, and the total number of alleles in called genotypes (AN) being AN>2,000. The ClinGen Sequence Variant Interpretation (SVI) group recommends that a dataset has a minimum of 2,000 alleles screened to utilize the AFs for variant interpretation^85^. We also applied a quality filtering on this WGS data requiring robust median coverage of ≥25X (Suppl. Fig. 1) for both the C and G bases comprising a CpG dinucleotide pair.

#### UK Biobank

UK Biobank (UKB) is a very large, population-based prospective study of 490,640 predominately European-ancestry individuals^84^. Characteristically, the genetic composition of these inidividuals is defined by five distinct ancestry cohorts: non-Finnish Europeans (n=458,855 – 93.52%), Africans (n=9,229 – 1.88%), Ashkenazi Jewish (n=2,869 – 0.58%), East Asians (n=2,245 – 0.46%), South Asians (n=9,674 – 1.97%) and remaining (n=7,768 – 1.58%). The WGS data for all individuals were released in December 2024 and accessed via the UKB DNA Nexus platform (https://ukbiobank.dnanexus.com/). We used the whole genome GraphTyper joint call pVCF files (Population level WGS variants) from the UKB. We applied quality filtering to the variant sites and filtered out variants with an AAScore<0.5 as recommended and employed previously^86,87^ and without a ‘PASS’ tag. The AAScore is a quality metric derived from a logistic regression model, ranging from 0 to 1, where higher values indicate greater probability of being a true positive variant. The PASS tag refers to the ratio of genotype calls that have passed QC.

### Identification of ultra-conserved CpGs (ucCpGs)

We screened the human genome to identify CpGs that were not observed to be mutated in these two datasets. CpGs were defined as ultra-conserved CpGs (ucCpGs) if neither cytosine (C) or gunanine (G) overlapped with a SNV and/or an Indel. Each CpG dinucleotide was considered as a single unit, and if either base was mutated, the site was not classified as an ucCpG. Using the bedtools (v2.31.1) intersectBed function^88^, ucCpGs were identified seperately within each dataset, and this resulted in two cohort-specific pre-quality control (pre-QC) non-polymorphic sets of ucCpGs. Quality control filters including minimum sequencing depth, AAScore and PASS status – were then applied to generate two robust, cohort-specific sets of ucCpGs. Finally, the two filtered ucCpG sets were intersected to determine the number of overlapping ucCpGs, which we define as the consensus set of ucCpGs used in all primary analyses.

#### Zoonomia PhyloP scores

To evaluate whether ucCpGs are more evolutionary conserved than vCpGs, we used base-wise conservation scores derived from the Zoonomia Project (https://zoonomiaproject.org/the-data/)^21^. The project alignment represents a total of 240 species and spanning ∼110 million years of mammalian evolution. For each CpG site, we computed the average PhyloP score across the two nucleotide positions to obtain a single conservation metric per CpG with negative scores representing acceleration and positive values conservation.

#### Genomic & Epigenomic Enrichment analyses

All statistical enrichment tests and displayed p values are derived from Fisher’s exact test performed via R (v4.5.1) unless specified.

#### Genomic Feature annotations in CpGs

All CpGs were annotated to genomic features using the ChIPseeker R package^89^. The CpG island (CGI) set was downloaded from UCSC https://hgdownload.soe.ucsc.edu/goldenPath/hg38/database/cpgIslandExt.txt.gz. CGI shores are ±2kb CGI surrounding and non-CGI overlapping regions.

#### ENCODE SCREEN

The ENCODE SCREEN (Search candidate cis-Regulatory Elements (cCREs)^25^ registry integrates DNase-seq, histone marks and transcription factor binding to classify high-confidence cCREs. SCREEN (v4) contains 1,063,878 human cCREs in GRCh38 (https://screen.encodeproject.org/) including promoters (PLS), proximal enhancers (pELS), distal enhancers (dELS) and CTCF-bound PLS, pELS and dELS.

#### Imprinting Control Regions

A list of 383 imprinting control regions (ICRs) was collated by Akbari et al. from four studies^28^. A ‘well-characterised’ ICR subset was primarily used comprising 68 ICRs with evidence from at least two studies.

#### Chromatin Segmentation

Chromatin segmentation analysis from the IHEC hg38-aligned EpiATLAS^90^ includes representative chromHMM CSREP 18-state data^91^. Annotations were downloaded from the pre-release portal for three representative healthy tissues, each corresponding to a distinct embryonic germ layer. Specifically, we selected hepatocyte (hepatocyteH_summary_18_ChromHMM.bed) as an endoderm-derived tissue, muscle (muscle_organH_summary_18_ChromHMM.bed) as a mesoderm-derived tissue and brain (brainH_summary_18_ChromHMM.bed) as an ectoderm-derived tissue.

#### Cell-type specific DNA methylomes

We downloaded the publicly available data from the DNA methylation atlas of normal human cell types from Loyfer *et al*.^27^. We used the top 1% of variable DNA methylation blocks results, defined as genomic regions of at least four CpG sites. The hg19 data were lifted to GRCh38 via the UCSC commandline tool liftOver. We also collected the information on a genome-wide set of unmethylated regions (UMRs) for each cell type across 39 tissues. UMRs had been defined as blocks containing at least four CpGs in which ≥85% of sequenced fragments were unmethylated in at least 85% of covered CpGs (∼.4CpGs.U85.f85 bed files). Cell type-specific UMRs, once lifted to GRCh38 were subsequently interrogated to delineate any potential tissue-specific enrichment of ucCpGs within these UMRs. A common pan-tissue set of UMRs - present in all 39 tissues - was also generated using bedtools multi-intersect and also lifted to GRCh38.

#### HUGO Gene Nomenclature Committee (HGNC) Gene Family Enrichment

The annotatr R package (v1.30.0)^92^ was used to classify all autosomal CpGs located within gene promoters, exons, 5’UTRs and 3’UTRs via Gencode v48 annotation. These annotated CpGs were then grouped according to HGNC gene families (downloaded on 31/05/25 from HGNC https://www.genenames.org/download). The ratio of ucCpGs to genomic CpGs was then calculated for these gene families. These were ranked by enrichment p value calculated via Fisher’s test and adjusted p-values were calculated via R p.adjust (p_value, method =“fdr”).

#### Human four-fold degenerate sites, human-specific CpGs, and Evolutionary Accelerated Regions

The set of ∼5.3M human four-fold degenerate sites within coding regions was downloaded from (https://doi.org/10.6084/m9.figshare.26318410)^45^. Datasets were also gathered on human-specific CpGs (https://zenodo.org/records/13377375)^48^, human-accelerated regions (HARs)^49^, human ancestor quickly evolved regions (HAQERs)^50^ and HAQER CpG beacons^51^.

#### Combined Annotation Dependent Deletion (CADD) Sequence scores & ClinVAR Pathogenic Variants

Combined Annotation Dependent Deletion (CADD) v1.7 data, including all possible SNVs of GRCh38 was downloaded (https://cadd.gs.washington.edu/download)^52^. The CADD Phred-like scores, range from 1 to 99, with larger values indicating more deleterious variants. These scores are based on the rank of each variant relative to all 8.6 billion possible substitutions in the human reference genome. To obtain a single deleteriousness metric per CpG site, we calculated the average CADD score across the two nucleotide positions.

ClinVAR pathogenic variants are curated from submission >2800 organizations across the world^53^. We downloaded the clinvar_20250824 VCF file from https://ftp.ncbi.nlm.nih.gov/pub/clinvar/vcf_GRCh38/. Variants were filtered to include only those with a clinical significance of ‘pathogenic’ for further analysis.

### Identification of Significant ucCpG clusters

#### 1kb window ucCpG Counts

To identify ucCpG clusters, we first used the bedtools (v2.31.1) slopBed function, to create 1kb windows around each ucCpG. Specifically, 499bp were added upstream and downstream of each ucCpG position and then the total number of ucCpGs was calculated within each of these 1kb ucCpG centred windows to give ucCpG/kb scores via the bedtools intersectBed -c count flag.

#### Mutability Likelihood of individual ucCpGs

To assess the likelihood of a CpG being ultra-conserved, we accounted for the inherent variability in mutation rates of CpGs^93^. As a result, we estimated the mutational likelihood of each CpG based on two key drivers: (i) the DNAm level of the CpG in the germline, given the well-known increase in mutability with increased methylation, and (ii) the surrounding base context considering the base before and the base after, as certain bases are associated with slightly lower mutation rates. To incorporate these effects, we incorporated both before and after trinucleotide context-specific mutation rates (Suppl. Table 1) to capture context-dependent mutational biases. We further adjusted for the effect of DNAm on mutation rates using methylation data from gnomAD, derived from germ cells across 14 developmental stages. These data included samples from pre-implantation embryos (sperm, oocyte, pronucleus, 2-cell. 4-cell, 8-cell, morula and blastocyst stage embryos) and primordial germ cells (7, 10, 11, 13, 17 and 19 week)^54^.

#### Permutation Assessment of ucCpG cluster Significance Threshold

To identity extreme ucCpG clusters that were unlikely to occur by chance alone, we performed a permutation analysis informed by mutational likelihood. Taking the developmental DNAm data discussed above and the mutation likelihoods calculated from both DNAm state and ±1bp flanking base context, we derived comparative mutation rates to weight the probability of CpG selection during permutation. The permutation was repeated 1,000 times, and for each randomly generated ucCpG set, we calculated the maximum ucCpG density window (ucCpGs/kb). The distribution of these maximum density values across 1,000 permutations was recorded to define the empirical significance threshold for the actual observed ucCpG clusters.

#### Sperm DNA methylation data

To further assess the robustness of our mutational likelihood permutation, we repeated this with data from an additional independent germline DNA methylation dataset. Specifically, we gained assess to the McGill Sperm Methylome Sequencing dataset, of pooled whole-genome bisulfite sequencing (WGBS) data from sperm DNA from 21 healthy normospermic male participants^55^. Data were requested through the European Genome-Phenome Archive (EGA; accession number EGAD00001004978) and were made available following approval from the study leads. These data were originally aligned to hg19 and were lifted to GRCh38 via liftOver (4,725 CpGs were lifted to two locations with the highest coverage location retained). CpG sites were categorised by methylation level as low (≤20%), intermediate (20–79%), or high (≥80%).

#### Common Structural Variants

To assess the overlap of identified ucCpG clusters with deletional structural variants (SVs), we used recent pangenome global diversity generated long-read sequencing data from Logsdon et al.^56^ We focused on these high quality common SVs, as earlier datasets containing all rare, including singleton variants, and some potentially false positive SVs cover a majority of the genome (*e.g.*, gnomAD SV v4.1 ∼68.9%). Furthermore, common deletional SVs are those more likely to be present in a homozygous state. To focus our analysis on regulatory regions that do not show an appreciable level of complete deletion in the human population, we filtered out any extreme ucCpG clusters that overlapped with these SVs.

#### TissueEnrich

The Tissue Enrich tool (https://tissueenrich.gdcb.iastate.edu/) was used to calculate tissue-specific gene enrichment for the ucCpG extreme cluster genes^57^. The input gene set was generated via GREAT (v4.0.4)^94^ with the nearest gene calculated via ±5kb basal no extension. Tissue-specific genes were defined by processing RNA-seq data from the Human Protein Atlas (HPA)^57^.

#### Gene Intolerance to Loss-of-function mutation

We categorised protein coding genes by their LOEUF (loss-of-function observed/expected upper bound fraction) score, a measure of a gene’s tolerance to mutation^59^. Scores were downloaded from UCSC Table Browser: assembly hg38; group: variation; track: gnomAD Constraint Metrics table: Gene LoF (pliByGene). Very low LOEUF values are indicative of essential genes for survival. We then compared those genes associated with ucCpG clusters (via GREAT ±5kb basal no extension) versus those without.

#### GREAT enrichment analysis

We applied the Genomic Regions Enrichment Annotations analysis software tool (GREAT v4.0.4)^94^ to the 298 extreme ucCpG clusters of potential functional significance. We performed the enrichment focused on potential local promoter settings (±5kb) as these clusters are CpG dense, i.e., no distal extension (GREAT settings: human genome assembly: hg38. Proximal: 5kb upstream and 5kb downstream, plus Distal: 0kb). Enrichment was assessed for Gene Ontology Biological Processes, Human phenotypes, and Mouse Single Knockout phenotypes. We ranked results by hypergeometric raw p values.

#### Transcription factor motif enrichment analyses

For TFBS motif enrichment, we used the MEME suite^95^ with datasets from HOCOMOCO Human^96^, JASPAR CORE vertebrates^97^ and Human Methylcytosine influenced TFs^58^. The H0mo sapiens C0mprehensive M0del C0llection (HOCOMOCO) v11 FULL collection provides 769 human TF binding models of A (high) -D (low) quality/confidence, including alternative motifs for single TFs. The JASPAR CORE (2022) vertebrates database contains 1,202 profiles derived from experimentally defined TF binding sites. The Human Methylcytosine collection includes 1,787 methyl-HT-SELEX and HT-SELEX motifs^58^.

#### Disease-related Genomics England Whole Genome Sequences

We additionally analysed WGS data from the disease-focused Genomics England 100,000 Genomes Project (100KGP)^98^. The latest available dataset includes the rare disease cohort of 62,698 samples (29,602 probands with rare disease and 33,096 of their parental/relative genomes). Individuals assigned to one of five broad super-populations represent the genetic composition of Genomics England samples: Europeans (79.5%), Africans (3.9%), South Asians (10.0%), East Asians (0.6%) and remaining (6.0%). Access to the genetic data (aggregate VCF files) for the 100KGP participants was provided through the Genomics England Research Environment (GeL RE), a secure cloud workspace after becoming a member and following project approval. For information on how to apply for access to Genomics England data, please see the website: https://www.genomicsengland.co.uk/research/data. We included variants that passed QC criteria and were annotated as PASS in the FILTER column. Whole genomes were aligned to the GRCh38 and had a mean sequencing depth of 32X (range, 27-54), with at least 95% of the reference genome at a depth greater than 15X.

We tested if there was a significant enrichment of variants in Rare Disease (RD) probands compared to RD parents/relatives using a one-sided hypergeometric test implemented in R (v4.5.1) using the phyper function. The population size (N) was defined as the total number of individuals in the RD population, with the number of successes in the population (M) corresponding to the total number of variant CpGs in the RD cohort. The sample size (n) was the number of RD probands, and the observed number of successes (k) was the number of mutated CpGs observed in RD probands.

### Ageing-related DNA methylation Datasets

#### DNA methylation Clock-related datasets

Details of the CpGs included in ageing-clocks were downloaded from the specific publications supplementary datasets: Horvath^62^; Hannum et al.^63^; Horvath SkinBlood^64^; Zhang et al.^65^; DunedinPACE^67^; Mitotic clocks: EpiTOC^68^, EpiTOC2 and hypo-Clock^69^. The entire set of ∼3.7M hg38 solo-WCGW CpGs^14^ was downloaded from https://zwdzwd.github.io/pmd. A Bonferroni threshold p<3.57×10^−3^ to account for the multiple clock analyses was employed.

#### Chronological age (cAge)

Chronological (cAge) results from Bernabeu et al.^16^ were downloaded from their Suppl. Data Additional file 4 (13073_2023_1161_MOESM4_ESM.xlsx). This contains Table S8 that includes the linear (2,275) and quadratic (56) CpG cAge predictors and their coefficients. Survival EWAS data were available at https://datashare.ed.ac.uk/handle/10283/4781.

#### Ageing Differentially Methylated Regions (age-DMRs)

Age-related DMRs (398) were derived from 3,001 peripheral blood MeDIP-seq whole genome methylomes (autosomal) from Acton *et al*.^71^, which identified that a subset of these age-DMRs were enriched for tRNA loci compared to their background likelihood (see publication for details). Briefly, linear models were fitted to age using the MeDIP-seq DNAm data, as quantile normalised RPM scores at each 500 bp window and these results are the implemented model with sequencing batch and major blood cell counts (lymphocytes, monocytes, neutrophils, and eosinophil cell-counts from haematological data) run as fixed effects [UCSC trackhub: http://epigenome.soton.ac.uk/TrackHub/hub.txt] and extended from the earlier focused analyses of this same dataset within GWAS LD Block regions (∼22.1% of genome)^73^ (See Suppl. Table 2 for age-DMR hg38 bedfile). These sequencing-derived DNAm score data in 500bp semi-overlapped windows are available at the EMBL-EBI EGA under Dataset Accession number EGAD00010000983, access is subject to request and approval by their Data Access Committee.

#### Age-DMR Permutation analyses

To assess the significance of the enrichment of ucCpGs within the ageDMRs, two 1,000x permutation analyses were performed to account for CpG density and CpG context effects. Firstly, random age-DMR sets (398) that matched the observed size and precise CpG density were selected from the autosomes and their ucCpG count was calculated. These 1,000 permutated ucCpG count results were then compared with the observed ucCpG count with age-DMRs. Secondly, random age-DMRs (398) were selected that matched the observed CpG context of the majority overlap (>50%) of each individual observed age-DMR, i.e., CpG island (79), shore (148), shelf (16), and ocean (155). With the ocean category this was further subdivided and matched to overlap with protein coding gene exons (8), introns (88), non-protein coding gene exon (10) and introns (30), and intergenic (19) (via Gencode v48 annotation). The ucCpG count for each random set of CpG context and size matched DMRs (345: 500bp; 49: 1kb; and 4: 1.5kb) was then calculated. The ucCpG/CpG ratio was then assessed for these 1,000 permutations and compared to the observed age-DMR ratio value. For description the top Age-DMRs were collocated to the nearest gene ±10kb via GREAT (v4.0.4).

#### Smoking metaEWAS

Tobacco-smoking EWAS results were derived from a meta-analysis from Hoang et al.^70^ (65,857 smoking-associated Differentially Methylated Cytosines: DMCs) and then converted to build hg38 via liftOver (3 results unlifted). ucCpGs for these were intersected (via BedTools) with these ∼65.8k smoking associated CpGs (FDR <0.05) and this was conservatively compared to the entire EPICv1 array background set, as the reduced post-QC CpGs were not available.

## Data and Code Availability

Results of this analysis are available in Suppl. Tables 1-25. Suppl. Data File 1 is available at (http://doi.org/10.5281/zenodo.18683384). Code for all analysis available at GitHub at https://github.com/epigenomemed/ucCpG_paper on publication.

UK Biobank (http://www.ukbiobank.ac.uk/) data are available upon application and with permission of UKB’s Research Ethics Committee.

The gnomAD 4.1.0, dataset is available for download at http://gnomad.broadinstitute.org. There are no restrictions on the aggregate data released or results derived from gnomAD data.

Data from the National Genomic Research Library (NGRL) used in this research are available within the secure Genomics England Research Environment. Access to NGRL data is restricted to adhere to consent requirements and protect participant privacy. Data used in this research include: aggregated combined.vcf (/gel_data_resources/main_programme/aggregation/aggregate_gVCF_strelka/aggV2/). Access to NGRL data is provided to approved researchers who are members of the Genomics England Research Network, subject to institutional access agreements and research project approval under participant-led governance. For more information on data access, visit: https://www.genomicsengland.co.uk/research.

## Acknowledgements

The project was supported by a Longevity Impetus Grant and Hevolution Foundation.

This research has been conducted using the UK Biobank Resource under Application Number 96802 and uses data provided by patients and collected by the NHS as part of their care and support. We gratefully acknowledge the participants of the National Genomic Research Library (NGRL), (Genomics England (2024) https://doi.org/10.6084/m9.figshare.4530893), whose contributions made this research possible. Secure access to the NGRL under project ID 1211 was provided by Genomics England, which delivers the NGRL in partnership with NHS England, and is wholly owned by the UK Department of Health and Social Care. The NGRL contains participants’ health data collected by the NHS as part of their care, along with samples and data from their participation in research, for which fully informed consent has been obtained. This includes genomic and clinical data provided through the NHS Genomic Medicine Service, as well as data obtained through research studies, including the 100,000 Genomes Project and the Generation Study, both of which are delivered in partnership with the NHS, and from other research cohorts involving external collaborators.

This research utilised QMUL’s Apocrita HPC facility, supported by QMUL Research-IT (http://doi.org/10.5281/zenodo.438045).

## Author contributions

PC designed experiments, anaylsed data, and co-wrote the paper; CGB conceived the study, designed experiments, analysed data, supervised this work and co-wrote the paper. Both authors reviewed and approved the final manuscript.

## References

1. Li, E., Bestor, T.H. & Jaenisch, R. Targeted mutation of the DNA methyltransferase gene results in embryonic lethality. Cell 69, 915–26 (1992).

2. Okano, M., Bell, D.W., Haber, D.A. & Li, E. DNA methyltransferases Dnmt3a and Dnmt3b are essential for de novo methylation and mammalian development. Cell 99, 247–57 (1999).

3. Sutcliffe, J.S. et al. Deletions of a differentially methylated CpG island at the SNRPN gene define a putative imprinting control region. Nat Genet 8, 52–8 (1994).

4. Edwards, C.A. & Ferguson-Smith, A.C. Mechanisms regulating imprinted genes in clusters. Curr Opin Cell Biol 19, 281–9 (2007).

5. Bird, A. The dinucleotide CG as a genomic signalling module. Journal of molecular biology 409, 47–53 (2011).

6. Hodges, E. Sequencing in High Definition Drives a Changing Worldview of the Epigenome. Cold Spring Harb Perspect Med 9(2019).

7. Duncan, B.K. & Miller, J.H. Mutagenic deamination of cytosine residues in DNA. Nature 287, 560–1 (1980).

8. Adams, C.J. et al. Regularized sequence-context mutational trees capture variation in mutation rates across the human genome. PLOS Genetics 19, e1010807 (2023).

9. Tarkhov, A.E. et al. Nature of epigenetic aging from a single-cell perspective. Nat Aging 4, 854–870 (2024).

10. Lopez-Otin, C., Blasco, M.A., Partridge, L., Serrano, M. & Kroemer, G. Hallmarks of aging: An expanding universe. Cell (2022).

11. Tong, H. et al. Cell-type specific epigenetic clocks to quantify biological age at cell-type resolution. Aging (Albany NY) 16, 13452–13504 (2024).

12. Teschendorff, A.E. et al. Age-dependent DNA methylation of genes that are suppressed in stem cells is a hallmark of cancer. Genome Res 20, 440–6 (2010).

13. Moqri, M. et al. PRC2-AgeIndex as a universal biomarker of aging and rejuvenation. Nat Commun 15, 5956 (2024).

14. Zhou, W. et al. DNA methylation loss in late-replicating domains is linked to mitotic cell division. Nat Genet 50, 591–602 (2018).

15. Teschendorff, A.E. & Horvath, S. Epigenetic ageing clocks: statistical methods and emerging computational challenges. Nat Rev Genet (2025).

16. Bernabeu, E. et al. Refining epigenetic prediction of chronological and biological age. Genome Medicine 15, 12 (2023).

17. Terekhova, M., Bohacova, P. & Artyomov, M.N. Human immune aging. Immunity (2025).

18. Lu, A.T. et al. Universal DNA methylation age across mammalian tissues. Nat Aging 3, 1144–1166 (2023).

19. Agarwal, I. & Przeworski, M. Mutation saturation for fitness effects at human CpG sites. Elife 10(2021).

20. Grimwood, J. et al. The DNA sequence and biology of human chromosome 19. Nature 428, 529–535 (2004).

21. Christmas, M.J. et al. Evolutionary constraint and innovation across hundreds of placental mammals. Science 380, eabn3943 (2023).

22. De, S. & Babu, M.M. A time-invariant principle of genome evolution. Proc Natl Acad Sci U S A 107, 13004–9 (2010).

23. Cortes Guzman, M., Castellano, D., Serrano Colome, C., Seplyarskiy, V. & Weghorn, D. Transcription start sites experience a high influx of heritable variants fueled by early development. Nat Commun 16, 10120 (2025).

24. Doi, A. et al. Differential methylation of tissue- and cancer-specific CpG island shores distinguishes human induced pluripotent stem cells, embryonic stem cells and fibroblasts. Nat Genet 41, 1350–3 (2009).

25. Abascal, F. et al. Expanded encyclopaedias of DNA elements in the human and mouse genomes. Nature 583, 699–710 (2020).

26. Stadler, M.B. et al. DNA-binding factors shape the mouse methylome at distal regulatory regions. Nature 480, 490–5 (2011).

27. Loyfer, N. et al. A DNA methylation atlas of normal human cell types. Nature (2023).

28. Akbari, V. et al. Genome-wide detection of imprinted differentially methylated regions using nanopore sequencing. eLife 11, e77898 (2022).

29. Tanay, A., O’Donnell, A.H., Damelin, M. & Bestor, T.H. Hyperconserved CpG domains underlie Polycomb-binding sites. Proc Natl Acad Sci U S A 104, 5521–6 (2007).

30. Mas-Ponte, D. & Supek, F. Mutation rate heterogeneity at the sub-gene scale due to local DNA hypomethylation. Nucleic Acids Research 52, 4393–4408 (2024).

31. Liu, Z. et al. Nr2f1 enhancers have distinct functions in controlling Nr2f1 expression during cortical development. Proc Natl Acad Sci U S A 121, e2402368121 (2024).

32. Kim, S. et al. DNA-guided transcription factor cooperativity shapes face and limb mesenchyme. Cell 187, 692–711 e26 (2024).

33. Nguyen, T.T. et al. TFAP2 paralogs regulate midfacial development in part through a conserved ALX genetic pathway. Development 151(2024).

34. Mannens, C.C.A. et al. Chromatin accessibility during human first-trimester neurodevelopment. Nature 647, 179–186 (2025).

35. Zheng, S. & Sheng, R. The emerging understanding of Frizzled receptors. FEBS Lett 598, 1939–1954 (2024).

36. Tycko, J. et al. High-Throughput Discovery and Characterization of Human Transcriptional Effectors. Cell 183, 2020–2035.e16 (2020).

37. Guo, Y. et al. Histone H2A ubiquitination resulting from Brap loss of function connects multiple aging hallmarks and accelerates neurodegeneration. iScience 25, 104519 (2022).

38. Belotti, E. et al. H2A.Z is involved in premature aging and DSB repair initiation in muscle fibers. Nucleic Acids Res 52, 3031–3049 (2024).

39. Stefanelli, G. et al. Learning and Age-Related Changes in Genome-wide H2A.Z Binding in the Mouse Hippocampus. Cell Reports 22, 1124–1131 (2018).

40. Contrepois, K. et al. Histone variant H2A.J accumulates in senescent cells and promotes inflammatory gene expression. Nature Communications 8, 14995 (2017).

41. Aguilar, G.R. et al. Functional analysis of conserved C. elegans bHLH family members uncovers lifespan control by a peptidergic hub neuron. PLOS Biology 23, e3002979 (2025).

42. Hu, H. et al. ZKSCAN3 counteracts cellular senescence by stabilizing heterochromatin. Nucleic Acids Research 48, 6001–6018 (2020).

43. Serrano-Saiz, E., Leyva-Díaz, E., De La Cruz, E. & Hobert, O. BRN3-type POU Homeobox Genes Maintain the Identity of Mature Postmitotic Neurons in Nematodes and Mice. Current Biology 28, 2813–2823.e2 (2018).

44. Kõks, S. et al. Mouse models of ageing and their relevance to disease. Mechanisms of Ageing and Development 160, 41–53 (2016).

45. Christmas, M.J., Dong, M.X., Meadows, J.R.S., Kozyrev, S.V. & Lindblad-Toh, K. Interpreting mammalian synonymous site conservation in light of the unwanted transcript hypothesis. Nat Commun 16, 2007 (2025).

46. Neri, F. et al. Intragenic DNA methylation prevents spurious transcription initiation. Nature 543, 72–77 (2017).

47. Branciamore, S., Chen, Z.X., Riggs, A.D. & Rodin, S.N. CpG island clusters and pro-epigenetic selection for CpGs in protein-coding exons of HOX and other transcription factors. Proc Natl Acad Sci U S A (2010).

48. Bell, C.G. et al. Human-specific CpG “beacons” identify loci associated with human-specific traits and disease. Epigenetics 7, 1188–99 (2012).

49. Doan, R.N. et al. Mutations in Human Accelerated Regions Disrupt Cognition and Social Behavior. Cell 167, 341–354.e12 (2016).

50. Mangan, R.J. et al. Adaptive sequence divergence forged new neurodevelopmental enhancers in humans. Cell 185, 4587–4603 e23 (2022).

51. Abeykoon, Y. et al. Interrogating the Regulatory Function of HAQERs during Human Cortical Development. bioRxiv, 2025.11.30.691411 (2025).

52. Schubach, M., Maass, T., Nazaretyan, L., Röner, S. & Kircher, M. CADD v1.7: using protein language models, regulatory CNNs and other nucleotide-level scores to improve genome-wide variant predictions. Nucleic Acids Res 52, D1143–d1154 (2024).

53. Landrum, Melissa J. et al. ClinVar: updates to support classifications of both germline and somatic variants. Nucleic Acids Research 53, D1313–D1321 (2024).

54. Chen, S. et al. A genomic mutational constraint map using variation in 76,156 human genomes. Nature 625, 92–100 (2024).

55. Chan, D. et al. Customized MethylC-Capture Sequencing to Evaluate Variation in the Human Sperm DNA Methylome Representative of Altered Folate Metabolism. Environ Health Perspect 127, 87002 (2019).

56. Logsdon, G.A. et al. Complex genetic variation in nearly complete human genomes. Nature (2025).

57. Jain, A. & Tuteja, G. TissueEnrich: Tissue-specific gene enrichment analysis. Bioinformatics 35, 1966–1967 (2018).

58. Yin, Y. et al. Impact of cytosine methylation on DNA binding specificities of human transcription factors. Science 356(2017).

59. Karczewski, K.J. et al. The mutational constraint spectrum quantified from variation in 141,456 humans. Nature 581, 434–443 (2020).

60. Selleri, L. & Rijli, F.M. Shaping faces: genetic and epigenetic control of craniofacial morphogenesis. Nat Rev Genet 24, 610–626 (2023).

61. Haghani, A. et al. DNA methylation networks underlying mammalian traits. Science 381, eabq5693 (2023).

62. Horvath, S. DNA methylation age of human tissues and cell types. Genome Biol 14, R115 (2013).

63. Hannum, G. et al. Genome-wide methylation profiles reveal quantitative views of human aging rates. Mol Cell 49, 359–67 (2013).

64. Horvath, S. et al. Epigenetic clock for skin and blood cells applied to Hutchinson Gilford Progeria Syndrome and ex vivo studies. Aging (Albany NY) 10, 1758–1775 (2018).

65. Zhang, Q. et al. Improved precision of epigenetic clock estimates across tissues and its implication for biological ageing. Genome Med 11, 54 (2019).

66. Levine, M.E. et al. An epigenetic biomarker of aging for lifespan and healthspan. Aging (Albany NY) 10, 573–591 (2018).

67. Belsky, D.W. et al. DunedinPACE, a DNA methylation biomarker of the pace of aging. Elife 11(2022).

68. Yang, Z. et al. Correlation of an epigenetic mitotic clock with cancer risk. Genome Biol 17, 205 (2016).

69. Teschendorff, A.E. A comparison of epigenetic mitotic-like clocks for cancer risk prediction. Genome Medicine 12, 56 (2020).

70. Hoang, T.T. et al. Comprehensive evaluation of smoking exposures and their interactions on DNA methylation. EBioMedicine 100, 104956 (2024).

71. Acton, R.J. et al. The genomic loci of specific human tRNA genes exhibit ageing-related DNA hypermethylation. Nature Communications 12, 2655 (2021).

72. Gogler, A. et al. HSPA2 influences the differentiation and production of immunomodulatory mediators in human immortalized epidermal keratinocyte lines. Cell Death Dis 16, 344 (2025).

73. Bell, C.G. et al. Novel regional age-associated DNA methylation changes within human common disease-associated loci. Genome Biology 17, 193 (2016).

74. Grant, S.M. et al. Long-Read epigenetic clocks identify improved brain aging predictions. bioRxiv, 2025.09.30.679553 (2025).

75. Haghani, A., et al. EnsembleAge: enhancing epigenetic age assessment with a multi-clock framework. Geroscience (2025).

76. Jacques, M. et al. DNA Methylation Ageing Atlas Across 17 Human Tissues. bioRxiv, 2025.07.21.665830 (2025).

77. Xie, W. et al. Epigenomic analysis of multilineage differentiation of human embryonic stem cells. Cell 153, 1134–48 (2013).

78. Ma, Y., Budde, M.W., Zhu, J. & Elowitz, M.B. Tuning Methylation-Dependent Silencing Dynamics by Synthetic Modulation of CpG Density. ACS Synth Biol 12, 2536–2545 (2023).

79. Boukas, L., Bjornsson, H.T. & Hansen, K.D. Promoter CpG Density Predicts Downstream Gene Loss-of-Function Intolerance. The American Journal of Human Genetics 107, 487–498 (2020).

80. Wagner, W. Epigenetic networks coordinate DNA methylation across the genome. Molecular Therapy.

81. Liu, Y. et al. Identification of SENP7 and UTF1/VENTX as new loci influencing clustered protocadherin methylation across blood and brain using a genome-wide association study. Molecular Psychiatry (2025).

82. Han, A.L. et al. Diverse ancestral representation improves genetic intolerance metrics. Nat Commun 16, 2648 (2025).

83. Shabalin, A.A. et al. RaMWAS: fast methylome-wide association study pipeline for enrichment platforms. Bioinformatics 34, 2283–2285 (2018).

84. Carss, K. et al. Whole-genome sequencing of 490,640 UK Biobank participants. Nature 645, 692–701 (2025).

85. Ghosh, R. et al. Updated recommendation for the benign stand-alone ACMG/AMP criterion. Human Mutation 39, 1525–1530 (2018).

86. Halldorsson, B.V. et al. The sequences of 150,119 genomes in the UK Biobank. Nature 607, 732–740 (2022).

87. Hofmeister, R.J., Ribeiro, D.M., Rubinacci, S. & Delaneau, O. Accurate rare variant phasing of whole-genome and whole-exome sequencing data in the UK Biobank. Nat Genet 55, 1243–1249 (2023).

88. Quinlan, A.R. & Hall, I.M. BEDTools: a flexible suite of utilities for comparing genomic features. Bioinformatics 26, 841–2 (2010).

89. Yu, G., Wang, L.-G. & He, Q.-Y. ChIPseeker: an R/Bioconductor package for ChIP peak annotation, comparison and visualization. Bioinformatics 31, 2382–2383 (2015).

90. Raby, J., Frosi, G., White, F., Laperle, J. & Jacques, P.-É. Leveraging the largest harmonized epigenomic data collection for metadata prediction validated and augmented over 350,000 public epigenomic datasets. bioRxiv, 2025.09.04.670545 (2025).

91. Vu, H., Koch, Z., Fiziev, P. & Ernst, J. A framework for group-wise summarization and comparison of chromatin state annotations. Bioinformatics 39(2023).

92. Cavalcante, R.G. & Sartor, M.A. annotatr: genomic regions in context. Bioinformatics 33, 2381–2383 (2017).

93. Mugal, C.F. & Ellegren, H. Substitution rate variation at human CpG sites correlates with non-CpG divergence, methylation level and GC content. Genome Biology 12, R58 (2011).

94. McLean, C.Y. et al. GREAT improves functional interpretation of cis-regulatory regions. Nature biotechnology 28, 495–501 (2010).

95. Bailey, T.L. et al. MEME SUITE: tools for motif discovery and searching. Nucleic Acids Res 37, W202–8 (2009).

96. Vorontsov, I.E., et al. HOCOMOCO in 2024: a rebuild of the curated collection of binding models for human and mouse transcription factors. Nucleic Acids Research 52, D154–D163 (2023).

97. Castro-Mondragon, J.A., et al. JASPAR 2022: the 9th release of the open-access database of transcription factor binding profiles. Nucleic Acids Research 50, D165–D173 (2021).

98. Cipriani, V. et al. Rare disease gene association discovery in the 100,000 GenomesProject. Nature (2025).

