## Supplementary Figures for "Conserved Human CpG Dinucleotides Identify Enriched Ageing-related Signals in Developmental Pathways and the Brain"


1. Distribution of median per-base coverage on chromosome 21 in gnomAD
2. Density of CpGs versus ucCpGs per 1kb windows
3. ucCpG grouping using a machine learning approach
4. cAge Quadratic Coefficients in uCpGs compared to vCpGs
5. ucCpGs in ageDMRs – 1000x Permutations
6. *HSPA2* expression from the Human Protein Atlas
7. *HSPA2* genomic locus
8. *LHFPL4* genomic locus
9. *PCDHG* genomic locus

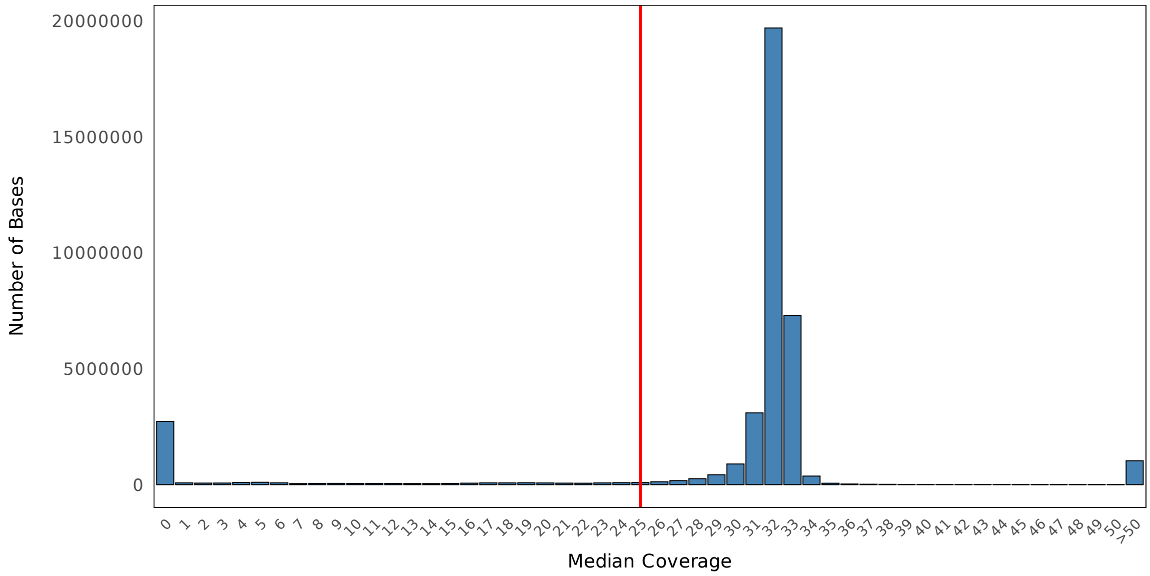


**Suppl Figure 1: Distribution of median per-base coverage on chromosome 21 in gnomAD**
Histogram showing the distribution of median per-base sequencing coverage across chromosome 21 in gnomAD whole-genome sequencing data. Coverage values above 50X are grouped for visualization clarity. The red vertical line indicates the 25X coverage threshold used to define reliably covered genomic regions.

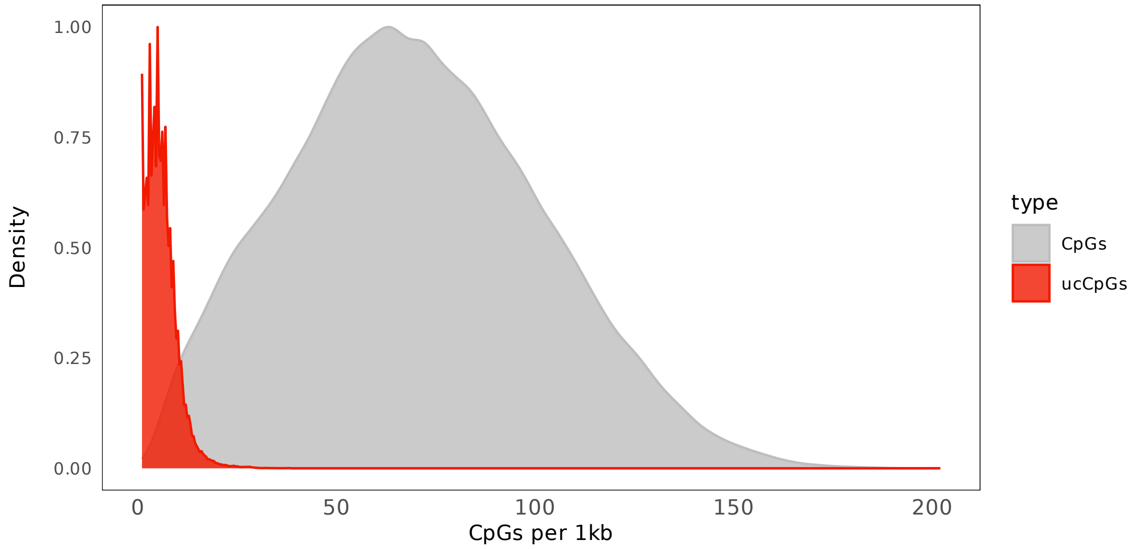


**Suppl Figure 2: Density of CpGs versus ucCpGs per 1kb windows**

The genomic CpG density (grey) across 1kb windows showed a unimodal distribution, with most windows containing ~47–92 CpGs/kb (~9–18% CpG of total sequence; CGIs generally >~12%, CpG shores ~5-12%) and a peak around 68–70 CpGs/kb (~14% CpG of total sequence). The ucCpG/1kb density was predominantly low (median:5, IQ range: 3-8).

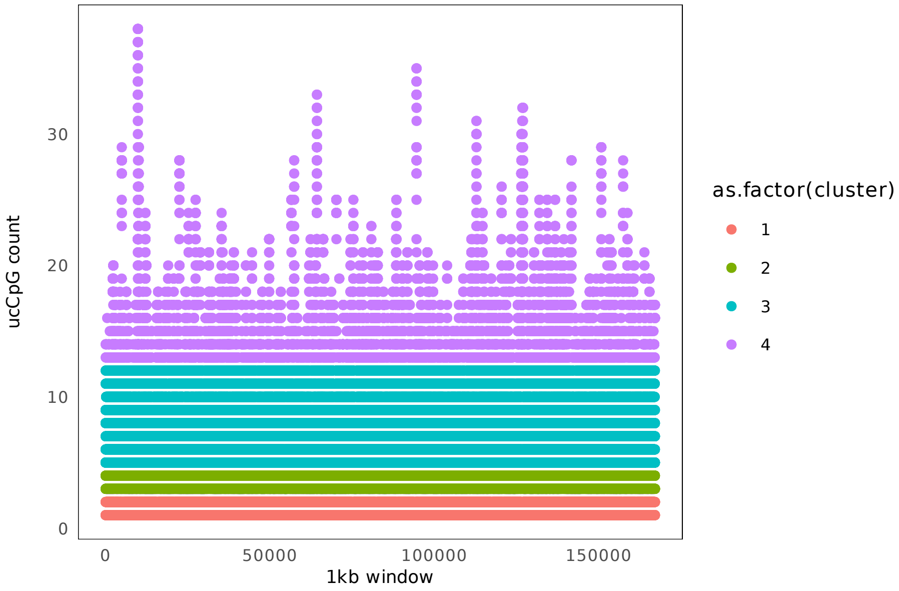


**Suppl Figure 3: ucCpG grouping using a machine learning approach**

An unsupervised machine learning approach using Gaussian mixture models (GMM) was applied to the ucCpG data. The model was fit on z-scored counts from ucCpG containing 1kb windows. In the 167,062 ucCpG windows, the GMM identified only 4 clusters for ucCpG windows. These corresponded to windows with 1-2 ucCpGs (Cluster1), 3-4 ucCpGs (Cluster2), 5-12 ucCpGs (Cluster3) and ≥13 ucCpGs (Cluster4).

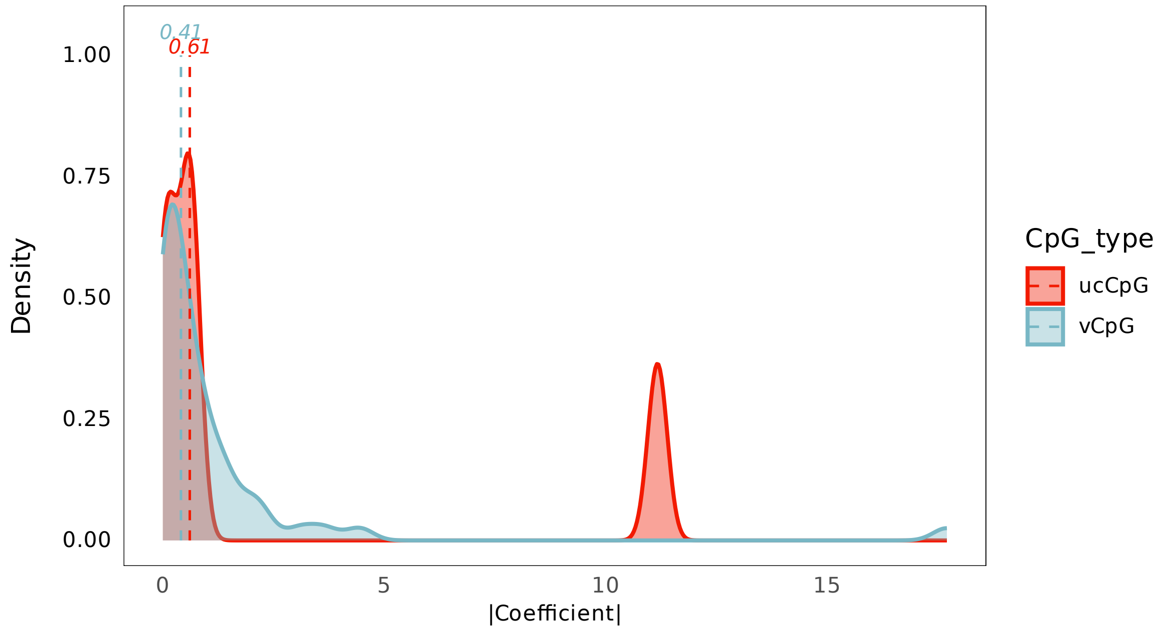


**Suppl Figure 4: cAge Quadratic Coefficients in ucCpGs compared to vCpGs**

Difference between absolute coefficients of ucCpGs and vCpGs for the chronological age (cAge) quadratic predictor CpGs from Bernabeu et al. (PMID:36855161). Median coefficient: ucCpG=0.61, vCpG=0.41. However, Wilcoxon test NS ($p>0.05$).

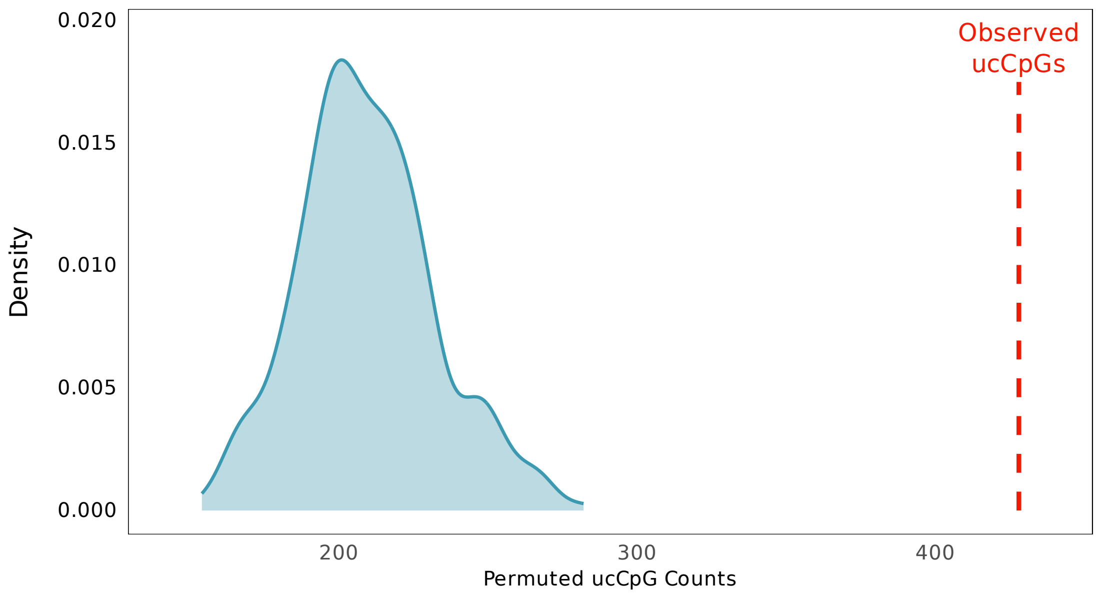


**Suppl Figure 5: ucCpGs in age-DMRs: 1000x Permutations**

Permutation by precise matched CpG density for each age-DMR (n=398). 1000x age-DMR permutations were performed for matched size (500bp=345; 1kb=49; 1.5kb=4) and CpG density. Then ucCpGs were counted in each of these random selections (min=139.0, 1st Q=193.0, median=207.0, mean=207.2, 3rd Q=221.0, max=287.0). No result reached the observed ucCpG age-DMR count of 428, Empirical $p<1x10^{-3}$.

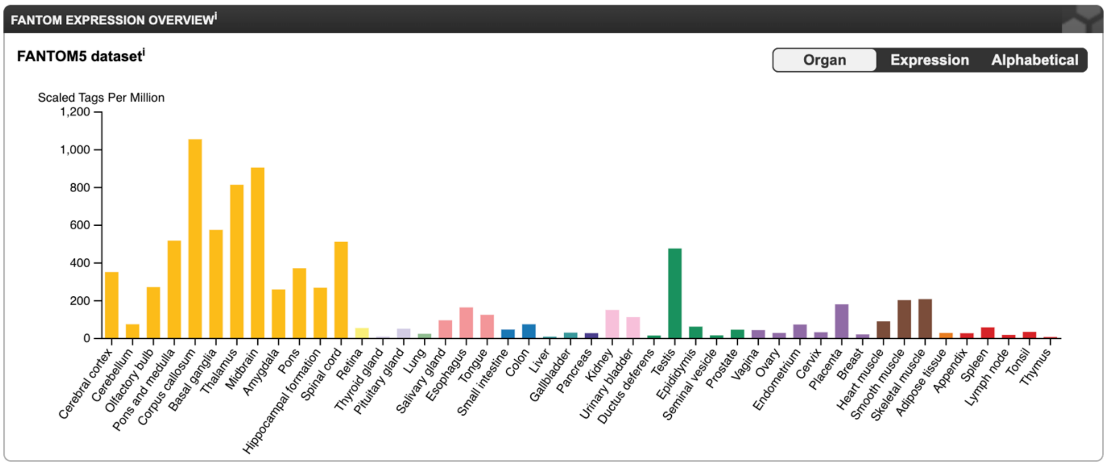


### Supp Fig 6: *HSPA2* Expression from the Human Protein Atlas

Expression levels across human tissues derived from FANTOM5 dataset.
https://www.proteinatlas.org/ENSG00000126803-HSPA2/tissue

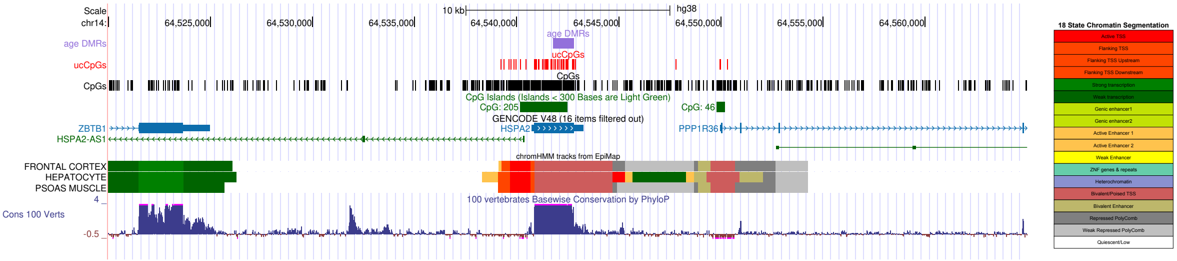


### Supp Figure 7: *HSPA2* genomic locus (hg38).

Tracks from top: Age-DMR (purple); ucCpGs (red); genomic CpGs (black); CpG islands (green); protein-coding genes (blue), non-coding genes (green), and pseudo-genes (magenta); and 100 vertebrates basewise conservation by PhyloP.

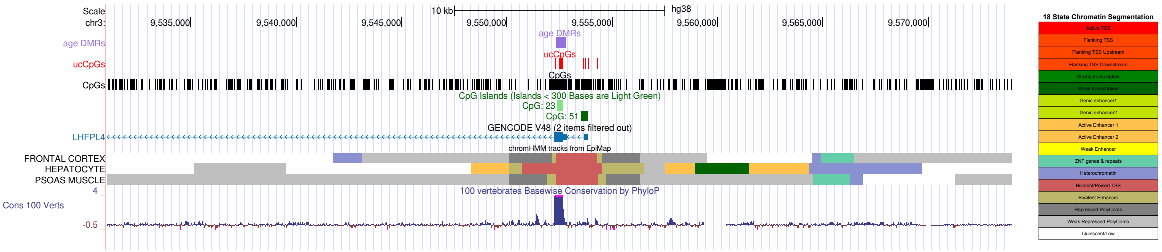


### Supp Figure 8: *LHFPL4* genomic locus (hg38).

Tracks from top: Age-DMR (purple); ucCpGs (red); genomic CpGs (black); CpG islands (green); protein-coding genes (blue), non-coding genes (green), and pseudo-genes (magenta); and 100 vertebrates basewise conservation by PhyloP.

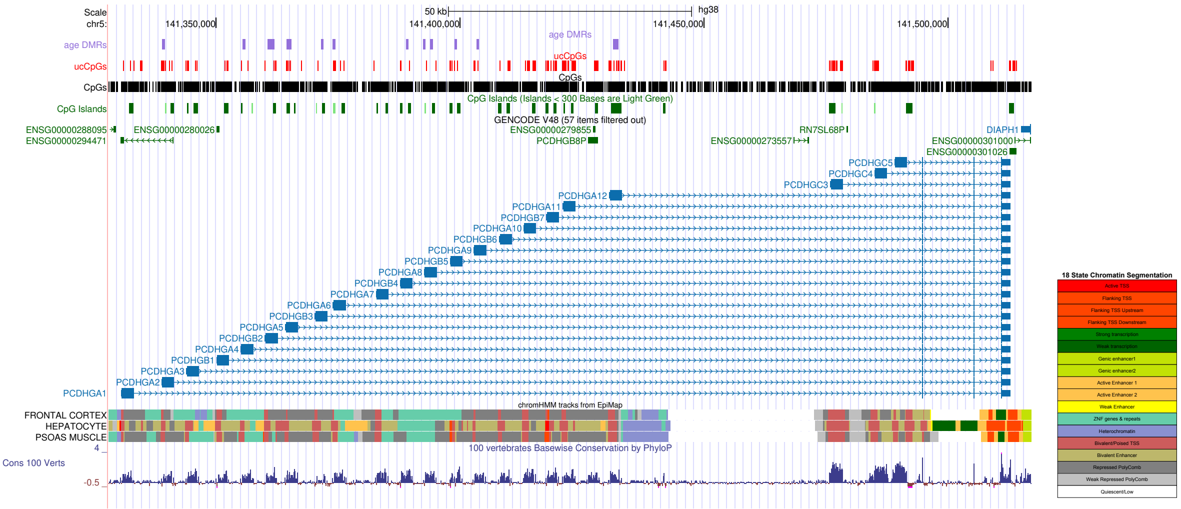


### Supp Figure 9: *PCDHG* genomic region (hg38)

Tracks from top: Age-DMR (purple); ucCpGs (red); genomic CpGs (black); CpG islands (green); protein-coding genes (blue), non-coding genes (green), and pseudo-genes (magenta); and 100 vertebrates basewise conservation by PhyloP.
